# Assessing pH-dependent Conformational Changes in the Fusion Peptide Proximal Region of the SARS-CoV-2 spike glycoprotein

**DOI:** 10.1101/2024.05.15.594386

**Authors:** Darya Stepanenko, Yuzhang Wang, Carlos Simmerling

## Abstract

One of the entry mechanisms of the SARS-CoV-2 coronavirus into host cells involves endosomal acidification. It has been proposed that under acidic conditions the Fusion Peptide Proximal Region (FPPR) of the SARS-CoV-2 spike glycoprotein acts as a pH-dependent switch, modulating immune response accessibility by influencing the positioning of the Receptor Binding Domain (RBD). This would provide an indirect coupling of RBD opening to environmental pH. Here, we explored this possible pH-dependent conformational equilibrium of the FPPR within the SARS-CoV-2 spike glycoprotein. We analyzed hundreds of experimentally determined spike structures from the Protein Data Bank, and carry out pH-Replica Exchange Molecular Dynamics, exploring the extent to which the FPPR conformation depends on pH and the positioning of the RBD. Meta-analysis of experimental structures identified alternate conformations of the FPPR among structures in which this flexible regions was resolved. However, the results did not support a correlation between the FPPR conformation and either RBD position or the reported pH of the cryo-EM experiment. We calculated pKa values for titratable side chains in the FPPR region using PDB structures, but these pKa values showed large differences between alternate PDB structures that otherwise adopt the same FPPR conformation type. This hampers comparison of pKa values in different FPPR conformations to rationalize a pH-dependent conformational change. We supplemented these PDB-based analyses with all-atom simulations, using constant pH-Replica Exchange Molecular Dynamics to estimate pKa values in the context of flexibility and explicit water. The resulting titration curves show good reproducibility between simulations, but also suggest that the titration curves of the different FPPR conformations are the same within error bars. In summary, we were unable to find evidence supporting the previously published hypothesis of FPPR pH-dependent equilibrium, either from existing experimental data, or from constant pH MD simulations. The study underscores the complexity of the spike system and opens avenues for further exploration into the interplay between pH and SARS-CoV-2 viral entry mechanisms.

## Introduction

The host cell entry mechanism of coronaviruses, such as the Severe Acute Respiratory Syndrome Coronavirus 2 (SARS-CoV-2), remains an active area of research. For SARS-CoV-2, the entry process is believed to initiate with the interaction between the viral spike fusion glycoprotein Receptor Binding Domain (RBD, shown in Figure 1) and the host cell receptor, angiotensin-converting enzyme 2 (ACE2). The RBD, which samples open and closed states, is able to interact with ACE2 only in the open state.^1,2^ Otherwise, the RBD receptor binding motif is not accessible for binding.^3^ Likewise, many antibody-binding epitopes on the RBD are masked in the closed state. Numerous structures are now present in the Protein Data Bank (PDB) with the three RBD domains in a variety of open and closed combinations.

**Figure 1:**
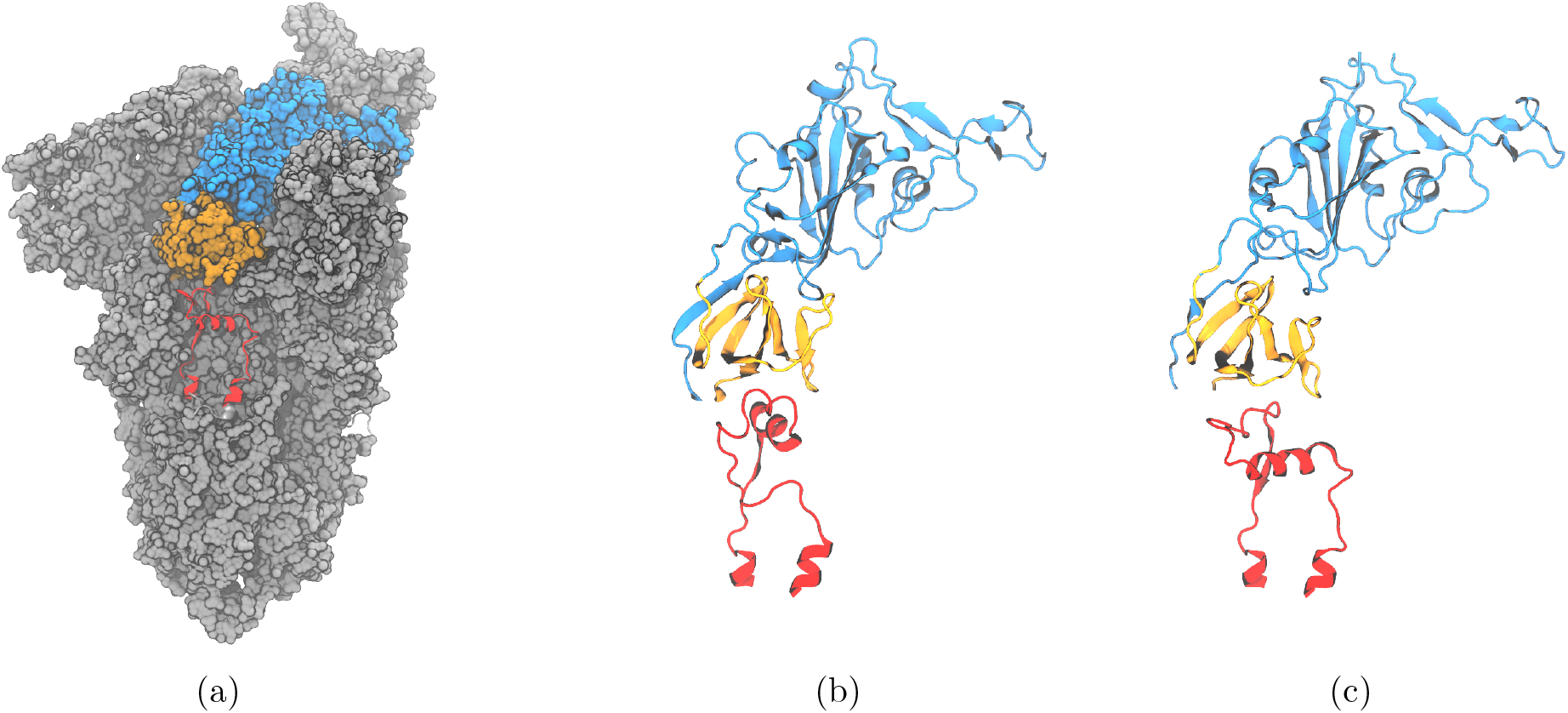
(a) Surface representation of spike trimeric ectodomain (PDB ID 6XM0). The Receptor Binding Domain is colored in blue, the C-terminal domain 1 in orange, and the Fusion Peptide Proximal Region in red; (b) Cartoon representation of RBD, CTD1 and compact conformation (PDB ID 6XR8 chain B^7^); (c) Cartoon representation of RBD, CTD1 and extended conformation FPPR (PDB ID 6XM0, FPPR from chain B^8^).

In one prior study of the SARS-CoV-2 2P (prefusion stabilized spike with K986P/V987P^4^), the authors observed only all-closed RBD conformation at pH 4.5 and 4.0.^5^ At pH 5.5, the authors observed RBD either in the open or closed position, or not in a defined position. This led the authors to hypothesize that acidification of the environment within the endosomal pH range, which varies from 6.5 to 4.0, mediates the positioning of the RBD. It was also proposed that this positioning allows the virus to evade the immune response within the endosome. Other authors^6^ have suggested that an all-closed spike structure at redduced pH may contribute to protein stability during viral assembly, preventing premature spike activation. In both cases, the authors suggest that pH-dependent RBD positioning may arise from pH-dependent conformational changes in a portion of the spike that is not in direct contact with the RBD; this region of the spike structure is discussed next.

### Fusion Peptide Proximal Region

The spike structure is shown in Figure 1. Directly under the RBD domain lies the C-Terminal Domain 1 (CTD1); under the CTD1 there is a short region which is unresolved in the majority of the structures deposited in the Protein Data Bank (PDB). This specific region, located between residues 824 and 858 with a disulfide bond between C840 and C851, is denoted as the **Fusion Peptide Proximal Region (FPPR)**. FPPR density was first resolved in the all-down-RBD configuration of the wild-type (WT) SARS-CoV-2 spike (PDB 6XR8).^9^ In that structure, the FPPR residues tightly pack together under the CTD1 and RBD of the protomer located in the clockwise direction from the FPPR, as illustrated in Figures 1(b) and 2(a). Hereafter, this conformation will be referred to as the **compact** FPPR.

**Figure 2:**
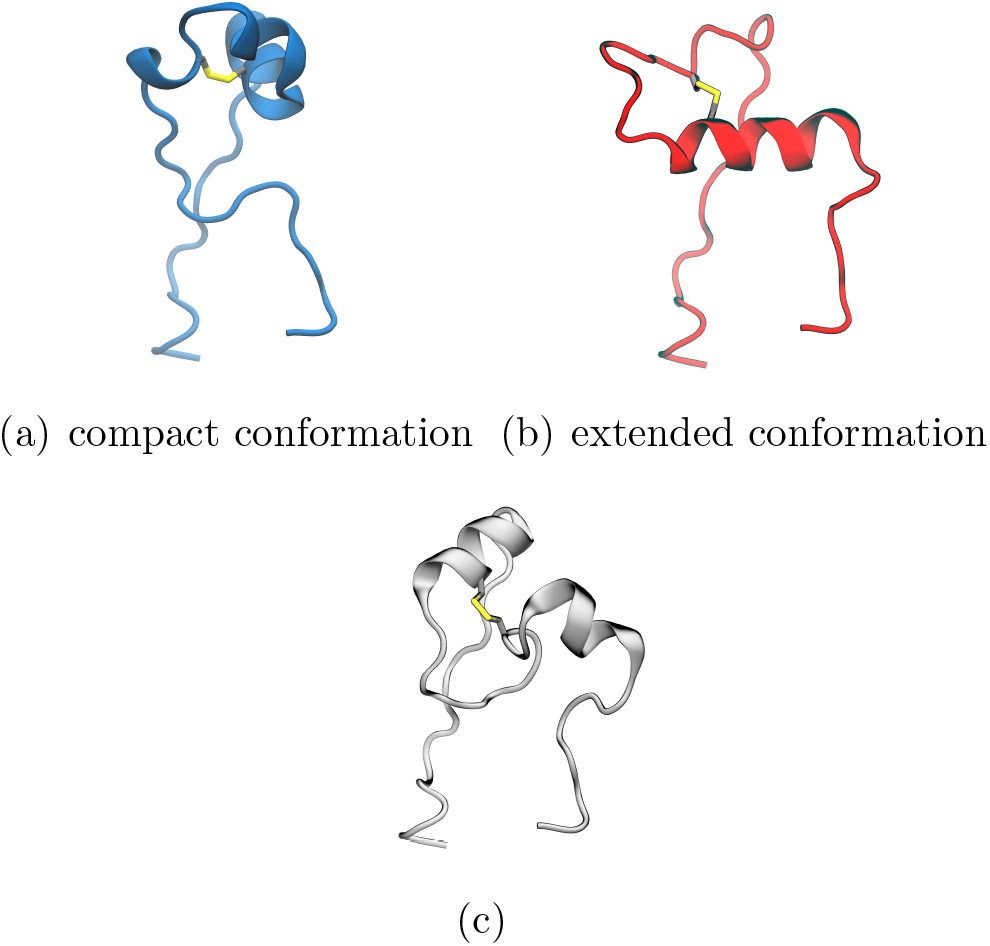
Representative structures for three conformational clusters of the FPPR obtained from analyzing spike structures in the PDB: **(a)** 7E7B (chain B), **(b)** 6XM0 (chain B), **(c)** 7LS9 (chain A)

Later, a different ordered FPPR conformation was reported (6XM0, using SARS-CoV-2 with the 2P mutation along with GSAS instead of RRAR at the S1/S2 furin cleavage site), also observed under a closed RBD, as shown in Figures 1(c) and 2(b).^5^ Here, this conformation will be referred to as the **extended** FPPR. The authors also reported a partially resolved, extended-conformation FPPR under an open RBD.

When the FPPR was initially resolved under a closed RBD,^9^ it was suggested that a structured FPPR (rather than the disordered FPPR as in all prior structures) might assist in maintaining the RBD in a downward position. The RBD and FPPR do not make direct contact, but are separated by the CTD1 domain. The authors observed that the CTD1 domain shifts downward when the RBD opens, which can cause a steric clash between CTD1 and a compact FPPR. Subsequently, upon the observation of both compact and extended conformations under closed RBDs, along with exclusively an extended FPPR (partially resolved) under open RBDs, it was proposed that the conformation adopted by the FPPR could influence the ability of the RBD to open, with the compact conformation aiding in keeping the RBD closed and only the extended or disordered conformation allowing RBD opening.^5^

The FPPR and surrounding region contain 13 acidic amino acids, including D614; the D614G mutation has largely replaced the original SARS-CoV-2 strain. These are partially compensated by nine nearby Arg or Lys. This high density of negative charge could lead to upward pKa shifts of these acidic side chains, possibly differing between alternate FPPR conformations in which the relative positions of the acidic groups vary. Some evidence exists to support a hypothesis that the FPPR conformation and pH are coupled. One cryo-EM spike study^5^ observed the compact FPPR conformation at pH values 4.0 and 5.5, but only the extended conformation at pH 5.5.^5^ Based on this observation, the authors suggested that FPPR serves as a pH-sensitive switch, with only the compact conformation being present under acidic conditions.^5^

Taken together, the two hypotheses (RBD position depends on FPPR, and FPPR is influenced by pH) imply that the RBD would be largely closed in an acidic pH environment, shedding bound ACE2 or antibodies^5^ and stabilizing the nascent spike during viral assembly.^6^

### Impact of acidic pH on spike behavior

There is additional evidence suggesting that the spike experiences conformational changes when exposed to acidic pH. It has been reported that acidic pH helps restore the spike protein after conformational changes that occur due to prolonged spike storage.^10^ Following the storage of the spike protein (SARS-CoV-2 2P variant) for 8 days at 4° C in PBS (pH 7.4), notable changes in conformational states were detected by negative-stain electron microscopy. The data indicate a reduction in the fraction of well-folded spike particles from 89% to 47%.^10^ Additionally, a decrease in the melting temperature was observed by differential scanning calorimetry (DSC). Upon 10 minutes exposure to buffers with pH 5.5 or 4.0, the DSC melting profile was restored, and a fully folded conformation was regained.^10^

Others have reported changes in spike stability with pH. After a disulfide-stabilized spike (S-R/x3, with a disulfide linking the RBD and central helix) was stored at 4° for 40 days at pH 7.4, most of the the Spike stayed folded, but the three RBDs moved slightly apart, and the FPPR became disordered, as observed by cryo-EM.^6^ Overnight incubation at pH 5.0 partially refolded the FPPR and brought the three RBDs closer.

In the present study, the hypothesis that the FPPR serves as a pH-sensitive switch was explored through meta-analysis of 700 experimental spike structures deposited in the PDB, and atomistic pH-Replica Exchange Molecular Dynamics simulations.

The analysis of PDB structures included extracting the RBD position, FPPR conformation, and reported pH. This revealed that both FPPR conformations have been observed in spike structures that were determined at pH ranging from 4 to 8; likewise, both conformations are observed under open and closed RBD domains, and both are present simultaneously in some spike structures. The lack of correlation between FPPR conformation and either pH or RBD position is supported by our constant pH simulations. We quantified FPPR dynamics at different pH values and estimated acidic side chain pKa values. The simulation results indicate that the net protonation of the titratable residues in the FPPR and surrounding amino acids is determined largely by the pH rather than FPPR conformation. These findings challenge the definition of the FPPR as a pH-sensitive switch, suggesting that both FPPR conformations may exist at neutral and acidic pH values, and factors other than pH may be responsible for observation of alternate FPPR conformations. Consequently, the role of the FPPR in spike protein dynamics and function remains unclear. We also propose that the extensive use of artificially stabilized spike constructs for structure determination may obscure the true biological behavior.

## Methods

All amino acid position numbers reported here are the same as in the full-length spike protein sequence.

700 SARS-CoV-2 full spike protein (Uniprot ID P0DTC2) cryo-EM structures with resolution 4 Å or better (see List S1) were retrieved from the PDB. The FPPR was identified using the sequence NKVTLADAGFI[KM]QYGDCLGD[IM]A[AY]RDLIC AQKF[NK]GLTVLPPLLTDEMI, where residues in [ ] indicate variants in the sequence. Next, structures were filtered to retain only those in which all alpha-carbon coordinates are present for the FPPR. Individual structures were categorized using two different labels: by the pH of the experiment as reported in the PDB file, or by the open/closed position of the RBD above that FPPR. RBD position was determined using the SpikeScape tool.^11^

### Clustering FPPR conformations from the PDB

First, FPPR coordinates were extracted from the full PDB files obtained using the filters described above. These FPPR conformations were clustered into different groups using the coordinates of the alpha carbon atoms. The hierarchical agglomerative (bottom-up) method as implemented in the *cpptraj*^*12*^ program was used, with a minimum distance between clusters of 4 Å.

### pKa values from static PDB structures

pKa values were calculated with PROPKA3.^13,14^

### Building the model system for MD simulations

Two independent systems were built, using experimental spike structures with compact and extended FPPR (6XLU and 6XM0). These were selected since they do not have any missing coordinates in or within 20 Å of the FPPR. Also, they were determined by the same lab,^5^ minimizing the possible impact of inconsistent structure determination protocols on our comparisons. The specific regions were: chain B, residues 35-54, 273-317, 726-783, 818-872 (the FPPR), 940-1023, 1054-1061 and chain A, residues 272-330, 530-60, 607-656, 661-673). To extract the coordinates of this region, the residues were selected in Pymol^15^ and then saved to a new PDB file. An example obtained using spike PDB ID 6XM0 is shown in Figure 3.

**Figure 3:**
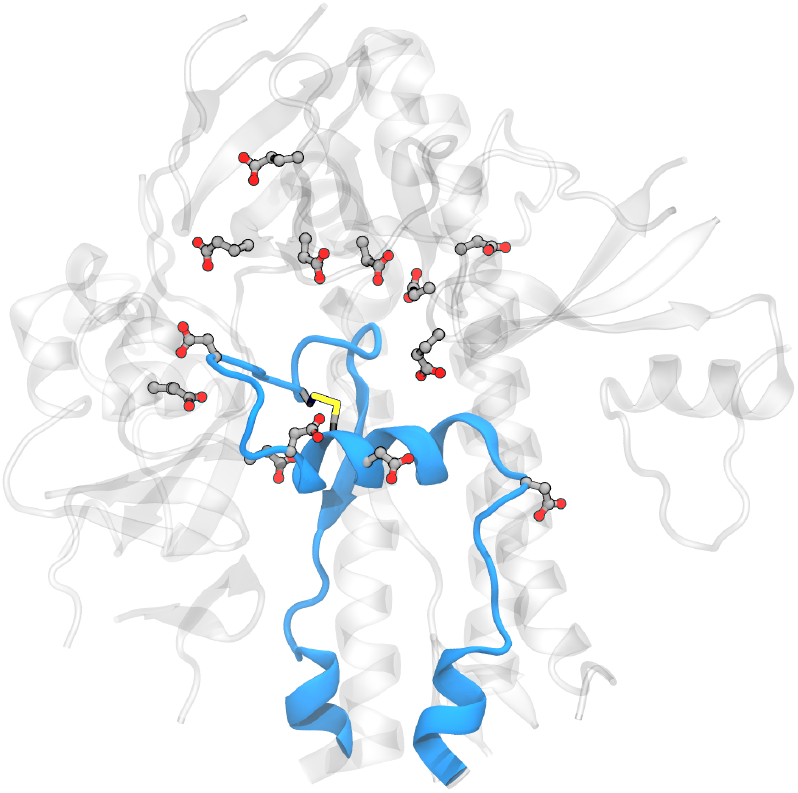
Model system for the spike FPPR region, showing the protein backbone in blue for the extended-conformation FPPR and gray for the surrounding region. Asp, Glu and Cys are shown in licorice. (PDB ID 6XM0, FPPR from chain B)

The 20 Å cutoff for the model system was selected based on the assumption that residues farther than 20 Å from the FPPR do not significantly contribute to the electrostatics of the titratable residues. The assumption was tested via calculations using PROPKA3, comparing pKa values obtained on the full spike as well as the smaller fragment obtained from the same PDB file. All pKa values of titratable residues in a fragment were within 0.5 units of those obtained from the complete spike structure, as shown in Table S1. This test provides an estimate of the pKa uncertainty introduced by using the smaller model.

### Molecular Dynamics General Protocol

The ff14SB^16^ and TIP3P^17^ force fields were used for the protein and water, respectively. Unless specified, all simulations used default settings in Amber v20^18^ with 2 fs time step at constant temperature and volume, an 8 Å cutoff with particle mesh Ewald^19^ for long range electrostatics, a Langevin thermostat with a collision frequency of 5*ps*^*−*1^, and SHAKE to constrain bonds involving hydrogens. Simulations were performed using pmemd.CUDA modules of Amber v20.^18^

For both systems, hydrogen atoms and water were added using the Amber *tleap* program, with a truncated octahedral periodic box with a minimum distance of solute to box edge of *∼* 15 Å ; the specific values varied in order to include the same number of water molecules in both systems (27734 molecules). 11 Na+ ions were added to neutralize the net charge of the system. The following residues were selected as titratable: Glu281, Asp830, Asp839, Asp843, Asp848, Glu554, Asp568, Asp571, Asp574, Glu583, Asp586, Asp614, Glu619 as shown in Figure 4. The final system consisted of 90405 atoms.

**Figure 4:**
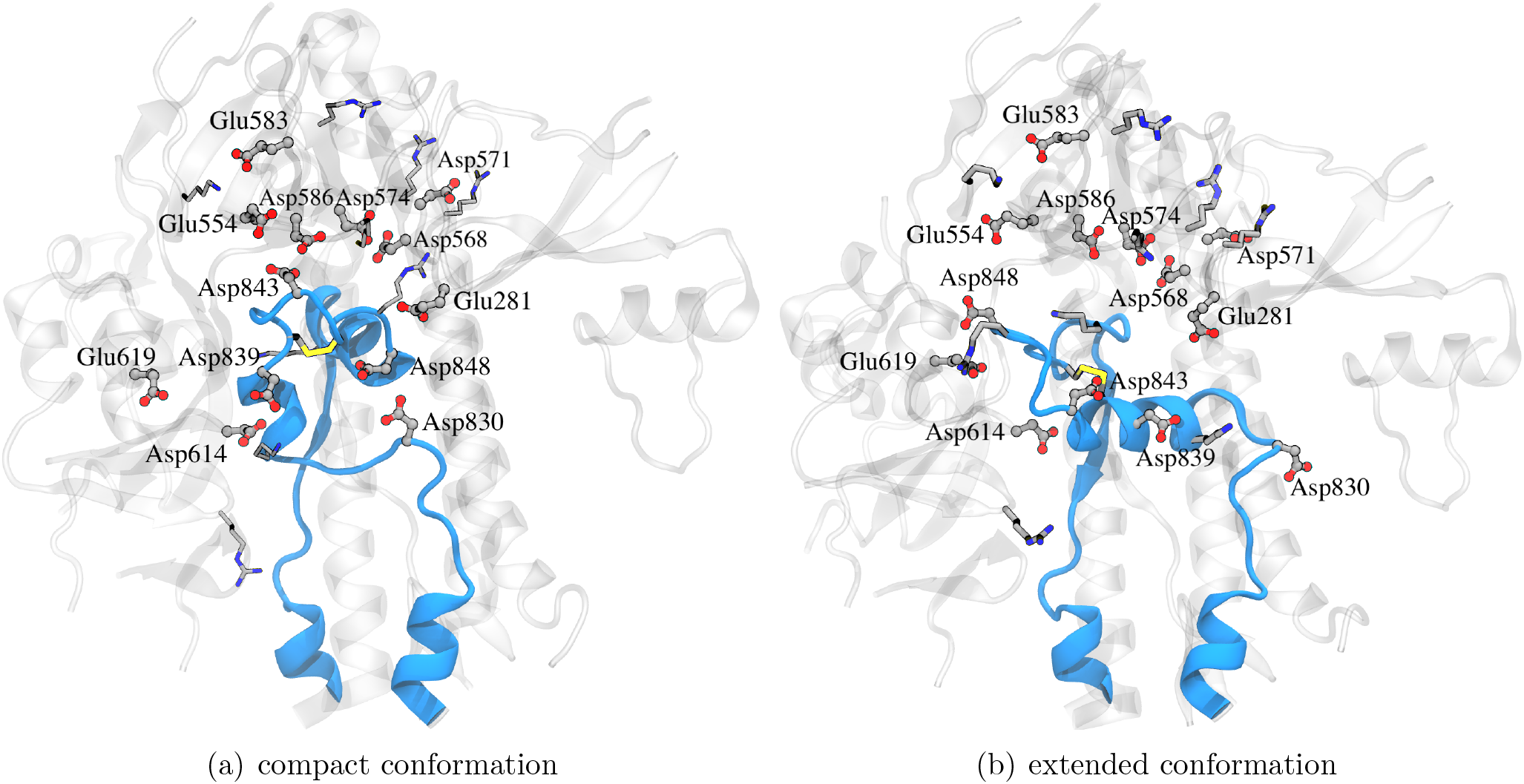
Part of spike glycoprotein with FPPR colored blue in (**a)** compact and **(b)** extended conformations with the rest of the protein around it colored gray with Asp, Glu, Lys and Arg in licorice. (PDB ID 6XLU, 6XM0, FPPR - chain B)

Both systems were relaxed with an 8 step protocol. Firstly, water and hydrogen atoms positions were minimized for 1000 steps using steepest descent and then for an additional 9000 steps with a conjugate gradient, while the rest of the system was restrained with 1 kcal/(mol Å^2^) Cartesian positional restraints.

Next, the system was heated to 300K at constant volume for 1 ns with 100 kcal/(mol Å^2^) restraints applied to everything except hydrogens and waters. The next steps involved 1 ns MD at constant pressure (1 bar) to relax the system density with the same positional restraints. Then, restraint force constants were lowered to 10 kcal/(mol Å^2^) for an additional 1 ns MD at constant pressure. Next, 10000 steps of conjugate gradient minimization was performed with restraints applied only to protein backbone atoms using a force constant of 10 kcal/(mol Å^2^). The next three steps of relaxation MD used 1 ns each at constant NPT, with positional restraints on protein backbone atoms with force constant of 10, 1, then 0.1 kcal/(mol*·*Å^2^). Each titratable carboxylate was deprotonated during relaxation, and no protonation state changes were attempted during these steps.

As only a subset of the spike protein was simulated, Cartesian positional restraints were maintained on all backbone atoms at the truncation boundary, along with backbone atoms of residues 272 to 291 in chain A and 292 to 317 in chain B, using a force constant of 10 kcal/(mol Å^2^)

### pH Replica Exchange Molecular Dynamics and estimation of pKa values

All pH-REMD simulations were performed with 8 replicas equally spaced 1 pH unit apart over the range of 1 to 8. The structure obtained after relaxation was used as initial coordinates and velocities for all replicas. A 2 fs time step was used and protonation state changes were attempted every 100 MD steps. For the protonation state change attempt, the Metropolis energy was calculated using the generalized Born implicit solvent model (igb=5 in Amber) with 0.1 M salt concentration.^20,21^ Intrinsic Born radii used the mbondi2 set,^20^ except for carboxylate oxygens in Asp and Glu which were changed to 1.3 Å to compensate for having 2 dummy protons present on each oxygen.^22,23^ An in-house, modified version of reference energies and modified cpinutil.py script were used, as the ones present in Amber are intended for usage with ff99SB force field. The reference energies were trained on model systems ACE-AS4-NME and ACEGL4-NME using ff14SB and TIP3P force fields, and were 31.578463 kcal/mol for the AS4 and 13.420729 kcal/mol for GL4. Force field files for Glu and Asp containing dummy protons (ASH and GLH) were adapted from the versions available for ff99SB in standard AmberTools 21. In frcmod.constph.ff14SB, AS4.ff14SB.off and GL4.ff14SB.off, the atom type of the C*β* atom was changed from CT to 2C, matching the change from ff99SB to ff14SB. Those files are available in SI.

During the constant pH REMD simulation, residues Glu281, Asp830, Asp839, Asp843, Asp848, Glu554, Asp568, Asp571, Asp574, Glu583, Asp586, Asp614, Glu619 were allowed to change protonation state. Attempts were performed every 100 MD steps. Each successful protonation change was followed by 100 steps of solvent relaxation.^22^

At every 1000 MD steps, exchanges were attempted between the solution pH values of neighboring replicas.

All ph-REMD simulations were run for 9 *·* 10^7^ steps or 180*ns* at constant volume. These simulations were performed in triplicate to assess reproducibility.

It was confirmed that protonation fractions of each of 13 titratable residues in both conformations were reasonably converged within 180 ns of simulation as illustrated in Figure S1 and S2. As illustrated in the pH-REMD pH ladder in Figure S3 and S4, structures from different replicas sampled multiple solution pH values throughout the simulation.

The pKa values of each of the 13 residues were calculated using the Levenberg-Marquardt algorithm from scipy.optimize.curve fit^24^ by fitting pH-REMD protonation fraction data at different pH values to the Hill equation as shown in Equation 1.

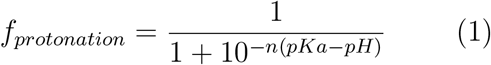

where *f*_*protonation*_ is a fraction of the total simulation that the titratable residue spent in a protonated state, and *n* is the Hill coefficient.

Backbone RMSDs and all atom symmetry-corrected non-hydrogen RMSDs were calculated with *cpptraj* .^12^ The clustering of pH-REMD simulations was performed using one of two methods mentioned in the text. The first method is a hierarchical agglomerative (bottom-up) approach. It considered the symmetrical RMSD of all non-hydrogen atoms with a minimum distance between clusters of 4 Å. This was used for the PDB analysis, where the expected number of clusters was not known. The second method used the k-means clustering algorithm on CA atoms. It had a setup of 2 clusters, a sieve of 3, and a randomized initial set of points^12^. This was used for the simulation analysis, where only two clusters were expected due to their use as initial coordinates. Visualizations were performed using VMD.^25^ The Python Matplotlib library was used for plot visualization.^26^

## Results and Discussion

### Experimental structural analysis

To test the hypothesis that the FPPR conformation is pH-dependent, all currently available SARS-CoV-2 trimeric ectodomain spike structures in the PDB (List S1) along with their reported experimental pH values were analyzed. To test the hypothesis that the compact FPPR conformation keeps the Receptor Binding Domain (RBD) closed, the positions of RBD domains above each FPPR were analyzed.

Out of the 700 analyzed spike structures, FPPR coordinates were present in 158 chains (see Methods for details). Structures in which the FPPR was not modeled were excluded from further analysis.

### Clustering PDB files

The 158 FPPRs were clustered using the alpha carbon coordinates, and 5 clusters were obtained. The distribution of structures among the five clusters is shown in Table 1 and Figure 5. The largest cluster contained 136 FPPRs, all of which resemble a compact conformation (the representative structure for this cluster is PDB ID 7E7B^27^), as shown in Figure 2(a). The second, smaller cluster consists of 12 FPPRs, and all structures in this cluster exhibited an extended conformation (representative structure for the cluster is PDB ID 6XM0^5^), as shown in Figure 2(b). The third cluster includes 6 structures and is illustrated in Figure 2(c) (representative structure is PDB ID 7LS9^28^). The fourth and fifth clusters include 3 and 1 structures, respectively; the PDB validation report of the representative structure of the fourth cluster (PDB 7N9T) states that the modeled FPPR does not fit well into the experimental density.^29–31^ while the single entry for the fifth cluster (PDB 7TPL) has a missing RBD above the FPPR.^32,33^

**Table 1:**
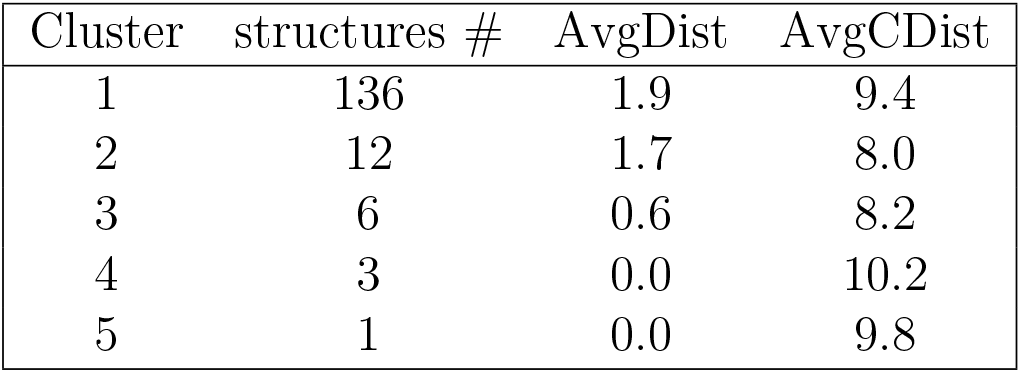
FPPR Cluster Statistics. AvgDist - average RMSD between all pairs of structures in the cluster, AvgCDist - average RMSD of this cluster to every other cluster.

| Cluster | structures # | AvgDist | AvgCDist |
| --- | --- | --- | --- |
| 1 | 136 | 1.9 | 9.4 |
| 2 | 12 | 1.7 | 8.0 |
| 3 | 6 | 0.6 | 8.2 |
| 4 | 3 | 0.0 | 10.2 |
| 5 | 1 | 0.0 | 9.8 |

**Figure 5:**
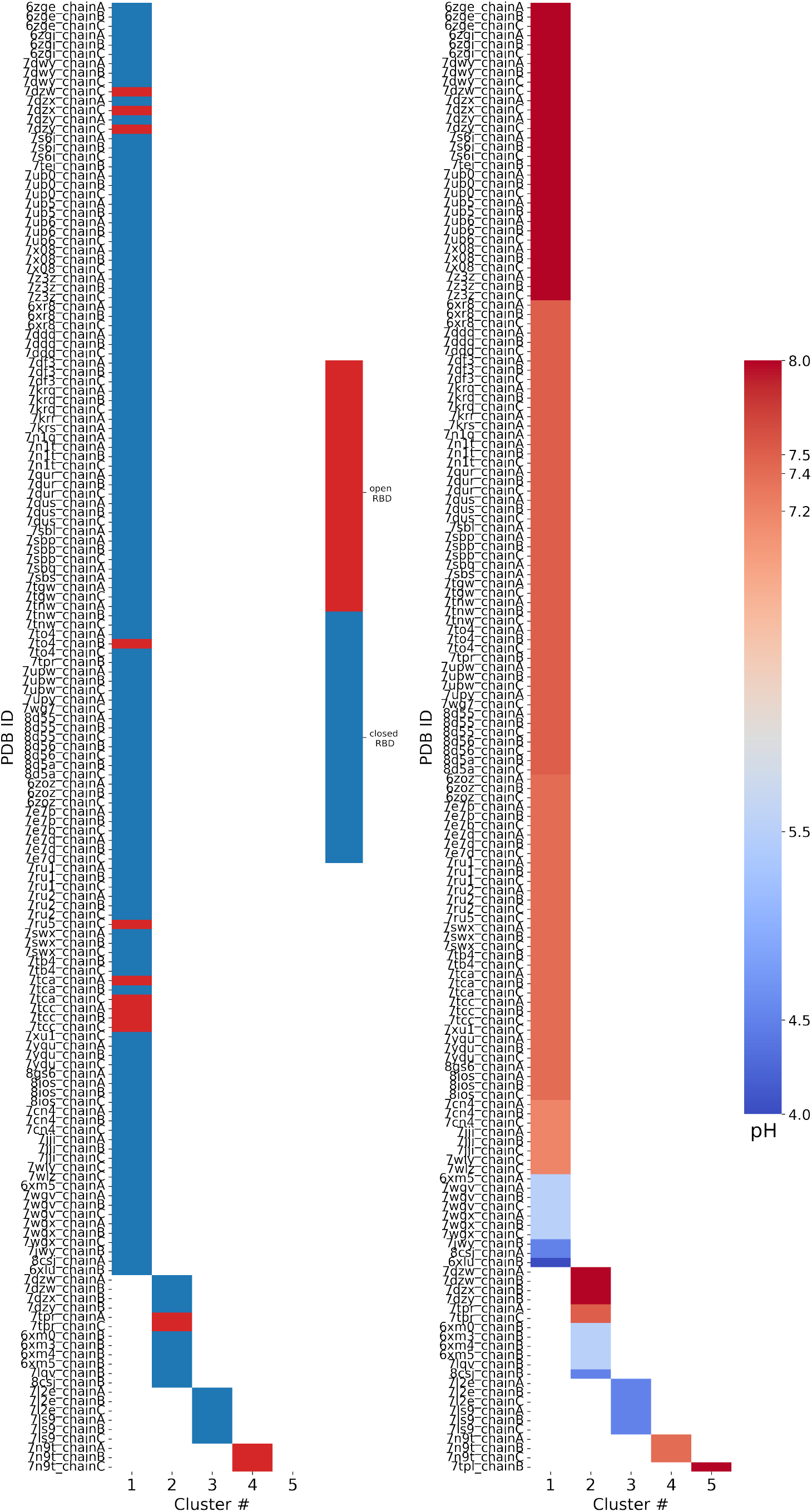
Results of FPPR cluster analysis applied to spike PDB structures. The PDB code and chain ID are given next to a colored box that is placed in the column according to the FPPR cluster number. The box color indicates the position of the RBD above the FPPR (left), or the pH of the experiment (right).

### RBD - FPPR conformation coupling

We analyzed spike RBD positioning using the SpikeScape^11^ tool. The majority of structures with a resolved FPPR have it positioned under the CTD1 connected to a closed RBD (142 of 158), as shown in Figure 5 (left). Among these closed-RBD structures, 126 have a compact FPPR, 10 have an extended FPPR, and 6 have a conformation from the 3rd cluster.

Among the 15 open-RBD structures with resolved FPPR, 10 have a compact FPPR, 2 have an extended FPPR, and 3 exhibit the FPPR conformation from the 4th cluster. The remaining FPPR structure has an unresolved RBD. Notably, three of the ten structures featuring the compact conformation modeled under an open RBD (7DZW,^34^ 7DZX^35^ and 7DZY^36,37^) exhibit steric clashes with the CTD1 region directly above the FPPR, as observed in the 3D structure (Figure S5) and PDB validation report.

With a closed RBD, the compact FPPR is more prevalent than the extended FPPR (126:10); under an open RBD, the preference for compact FPPR relaxes somewhat (7:2 after removing those with clashes). An alternate view is to focus on the FPPR; above a compact FPPR, the RBD closed:open ratio (126:7) is somewhat higher than for the extended FPPR (10:2).

Although a compact FPPR appears more frequently than extended FPPR when under a closed RBD, it remains that the compact FPPR conformation has been reported under an open RBD, casting doubt on the hypothesis that a compact FPPR conformation locks the RBD in a closed position. Moreover, among the open-RBD structures, the FPPR is more frequently modeled in the compact than extended conformation. However, the scarcity of structures with a fully resolved FPPR positioned under an open RBD complicates drawing reliable conclusions regarding a relationship between FPPR conformation and RBD position. We therefore shift focus to the role of pH.

### pH - FPPR conformation coupling

To explore the relationship between FPPR and pH, the reported pH during structure determination was extracted from the PDB files. As shown in Figure 5 (right) 136 of 158 structures report a neutral pH ranging from 7.2 to 8.0, while only 22 structures were resolved at pH values 4.0 to 5.5. However, the two most populated FPPR clusters, representing the compact and extended conformations, are both present across the entire pH range from 4.0 to 8.0.

Interestingly, there are cases at both neutral and acidic pH where the compact and extended FPPR conformations are present within the same spike structure (in different chains), despite the RBDs above these FPPRs all adopting the same open or closed position. Examples include 7DZX (pH 8), 7DZY (pH 8), 8CSJ^30^ (pH 4.5), and 6XM5^5^ (pH 5.5). The sampling of alternate FPPR conformations in the same spike structure suggests a small free energy difference between FPPR conformations, even at different pH values. These observations undermine the hypothesis that the compact conformation is strongly favored at acidic pH. However, as with the relatively rare open-RBD structures with ordered FPPR, the scarcity of spike structures determined at acidic pH makes it difficult to draw reliable conclusions.

In summary, analyzing all spike structures from the PDB yields results inconsistent with strong coupling between FPPR conformation and either experimental pH or RBD position, with both FPPR conformations observed in both neutral and acidic environments, as well as under both open and closed RBDs (sometimes in the same experimental structure). However, the reliability of the analysis is reduced by the scarcity of structures determined at acidic pH, or with the extended FPPR, along with issues related to poor-quality regions of these cryo-EM structure models. Given these uncertainties, we explored avenues to supplement the PDB-based analysis.

### Calculating pKa values for titratable side chains using PDB structures

If the equilibrium ratio of two protein conformations shows pH dependence, the titration curves of these two conformations should differ.^38^ Comparison of calculated pKa values for different conformations has been used to rationalize pH-dependent conformational changes.^39–43^ In the case of the FPPR, if the equilibrium preference between compact and extended FPPR has pH dependence, there should be at least one residue with different pKa values in those two conformations. The reference pKa values of Asp and Glu amino acids in water are 3.8 and 4.5. However, the pKa values in a protein could be quite different from those due to interactions with the local enviroment (e.g. formation of a salt bridge leads to a lower pKa). To test if the titration curves of compact and extended FPPRs are different, we calculated pKa values of titratable residues in the FPPR region for both conformations.

pKa values of titratable amino acids in the FPPR have been calculated previously.^5^ It was reported that the pKa values of Asp830, Asp843, Asp574, Asp586, and Asp614 in the compact conformation were higher than in the extended conformation, resulting in increased protonation of the FPPR region at pH 4.0. This result supports the hypothesis of pH-dependent FPPR conformation. However, the pKa calculations in that study were performed on static structures. Given the sensitivity of pKa values to local electrostatic interactions, it is possible that pKa values obtained using cryo-EM models may have larger uncertainties than those estimated using high-resolution crystal structures.

To assess the reproducibility of pKa calculations for the spike, pKa calculations were conducted here for multiple structures exhibiting each FPPR conformation. Specifically, two structures with the compact FPPR conformation and two structures with the extended FPPR conformation were selected, with the expectation that the differences in pKa values between spike structures exhibiting different FPPR conformations would be larger than those obtained from pairs of structures adopting the same FPPR conformation. For the extended conformation, coordinates from PDB entries 6XM0^5^ (determined at pH 5.5) and 7LQV^44^ (pH 5.5) were used, while for the compact conformation, 6XLU^5^ (pH 4) and 7WGX^45^ (pH 5.5) were used.

To measure the pKa values of titratable residues, the widely-used software PROPKA3^13,14^ was utilized. This software uses a physics-based model with empirical adjustments to calculate pKa values for a static structure. The resulting pKa values are presented in Table 2. For the extended FPPR conformation, pKa values from different spike structures agree within 1 pH unit. In contrast, the pKa values in the compact conformation for Glu583, Asp614, and Asp843 display differences of more than 1.5 units between two structures adopting the same FPPR conformation. Such discrepancies might be attributed to the local differences in structures. For example, in structure 6XLU, Asp843 is near Asp586 (see Figure S6), leading to stronger preference for the protonated acid and an increased pKa of 6.0. Conversely, in 7WGX where the carboxyl group is solvent-exposed, the pKa returns to 3.8.

**Table 2:** The pKa values from PROPKA3 for 13 titratable residues in example PDB structures adopting different FPPR conformations. Residues with a difference of 1.5 units or more in pKa values within the same conformation are colored red.

| Residue | FPPR Chain | Extended conformation |  | Compact conformation |  |
| --- | --- | --- | --- | --- | --- |
|  |  | 6XM0 | 7LQV | 6XLU | 7WGX |
| Glu 281 | B | 3.6 | 4.0 | 4.1 | 4.0 |
| Glu 554 | A | 4.2 | 3.9 | 4.6 | 3.9 |
| Asp 568 | A | 4.7 | 4.9 | 6.1 | 5.9 |
| Asp 571 | A | 4.3 | 4.9 | 4.8 | 5.1 |
| Asp 574 | A | 6.0 | 6.9 | 5.4 | 5.5 |
| Glu 583 | A | 4.9 | 5.3 | 4.9 | 3.4 |
| Asp 586 | A | 4.7 | 5.0 | 5.0 | 4.9 |
| Asp 614 | A | 5.5 | 5.8 | 7.2 | 4.9 |
| Glu 619 | A | 4.9 | 4.9 | 4.2 | 3.6 |
| Asp 830 | B | 4.2 | 4.6 | 4.5 | 4.1 |
| Asp 839 | B | 3.9 | 3.8 | 3.6 | 4.5 |
| Asp 843 | B | 4.2 | 4.3 | 6.0 | 3.8 |
| Asp 848 | B | 4.4 | 4.5 | 4.9 | 3.8 |

This analysis highlights how the specific choice of spike model from the PDB significantly affects pKa values even when the backbone conformation is highly similar, making the quantitative comparison of pKa values between different conformations unreliable.

## Constant pH REMD simulations

As demonstrated above, pKa calculations from static structures are sensitive to the details of the input experimental model. In contrast, molecular dynamics simulations are widely used to explore flexibility in biomolecular systems, sampling a dynamic ensemble in the context of an all-atom model with explicit water.^46^ We therefore explored whether atomistic molecular dynamics simulations could provide estimates of pKa values for acidic side chains in the FPPR region. MD simulations can sample local flexibility, which we hypothesized would lead to more precise pKa values. We also assess whether simulations are capable of directly sampling the interconversion between FPPR conformations, or show a preference for compact vs. extended FPPR under different pH conditions.

Traditional MD simulations employ a fixed protonation state for each titratable group that is set prior to the simulation (typically using the amino acid reference pKas, or less often, predictions from a static pKa calculation). Thus protonation states during these simulations will not be conformation-dependent.

A recent addition to the MD toolbox is constant pH molecular dynamics (cpHMD), which adds the ability to maintain a constant system pH rather than protonation state. CpHMD has been used to estimate pKa values with explicit inclusion of conformational dynamics, and to explore the coupling of protonation state and conformational equilibrium.^41,47,48^ In the discrete protonation variant of cpHMD that is used here, protonation state changes are attempted at intervals during the simulation, with the probability of accepting the change depending on the desired pH, the reference energy for the titratable amino acid, and the electrostatic energy of the system before and after the trial addition or removal of the proton.^21^ Performing cpHMD simulations over a range of pH values provides titration curves for each group, from which pKa values can be estimated. A more advanced version of the method, pH replica exchange MD (pH-REMD),^49–51^ allows simulations at different pH values to exchange conformations, leading to increased conformational sampling and better convergence. During pH-REMD, a series of coupled simulations (“replicas”) at different pH values are performed. At intervals, replicas can swap solution pH values, with the probability of accepting the swap based on the pH values and number of protons in each conformation. pH-REMD was used here to estimate pKa values for the FPPR region.

Achieving proper statistical significance requires extensive sampling in both structural and protonation spaces. However, the size of the trimeric spike in explicit water exceeds 1 million atoms,^52^ for which convergence of protonation states and conformational ensembles becomes intractable with current computational resources. Therefore, we opted to focus on a smaller model of the spike, including a single FPPR and any portion of the spike within 20 Å (see Methods).

Two simulation systems were built in explicit solvent, including the compact and extended FPPR. Thirteen amino acids were designated as titratable; these matched the ones examined for the static pKa calculations described above in Table 2. Eight equally spaced replicas, spanning pH values from 1 to 8 in unit increments, were generated. Although the pH of biological interest ranges from 4 to 7, the range from 1 to 8 was selected as a ladder for better sampling and titration curve fitting. Then, pH-REMD simulations of 180 ns were generated, using an initial structure with either the compact (PDB ID 6XLU^5^) or extended (6XM0^5^) FPPR conformation. For each conformation, pH-REMD simulations were repeated in triplicate to assess reproducibility of the calculated pKa values.

### No transitions between FPPR conformations are observed

Ideally, frequent transitions between the compact and extended FPPR conformations could be sampled, providing an estimate of the equilibrium constant for the FPPR conformational change at various pH values. However, no transitions between compact and extended conformations were sampled at any pH throughout the 180ns pH-REMD simulation. This was confirmed by comparing the FPPR backbone RMSD during time to both experimental conformations. The initially-extended FPPR simulation exhibited a low RMSD to the starting structure, with most backbone RMSD values ranging between 1 and 3 Å regardless of the pH (Figure 6 and S7). When compared to the compact FPPR, the RMSD for this initially-extended simulation was much larger, ranging between 6 Å and 8 Å as shown in Figure 7. This indicates that pH-REMD does not sample the compact FPPR conformation when started in the extended conformation. The initially-compact FPPR simulation exhibited more variability, with backbone RMSD values to the compact FPPR generally remaining below 4 Å but occasionally spiking up to 6 Å, as shown in Figure 6 and S7. However, no transitions to the extended FPPR were sampled, with RMSD values for the initially-compact simulations remaining above 6-8 Å. In summary, simulations starting from either FPPR conformation remained relatively stable at all pH values, and no transition to the other FPPR conformation was sampled. This slow rearrangement is consistent with the trapping of alternate FPPR conformations in some cryo-EM spike structures.

**Figure 6:**
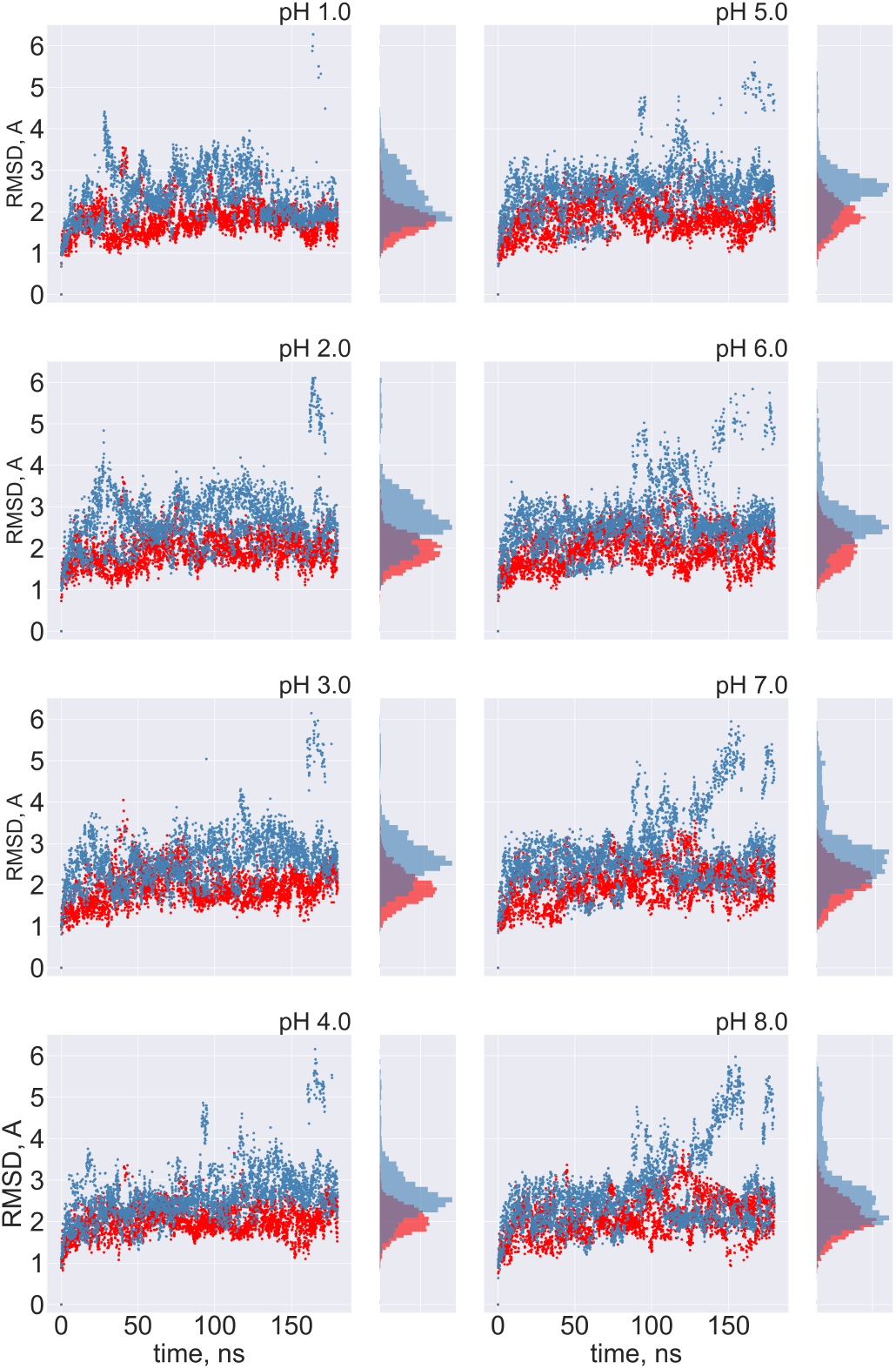
The FPPR remains in the intial conformation during simulations of extended and compact FPPR. FPPR RMSD values are shown at different pH values during two pH-REMD simulations, each initiated with a different FPPR conformation. Red symbols - initially extended simulation, with backbone RMSD calculated using the experimental extended conformation; blue symbols - initially compact simulation, with backbone RMSD calculated using to the experimental compact conformation. Low values indicate that the initial FPPR conformation is stable.

**Figure 7:**
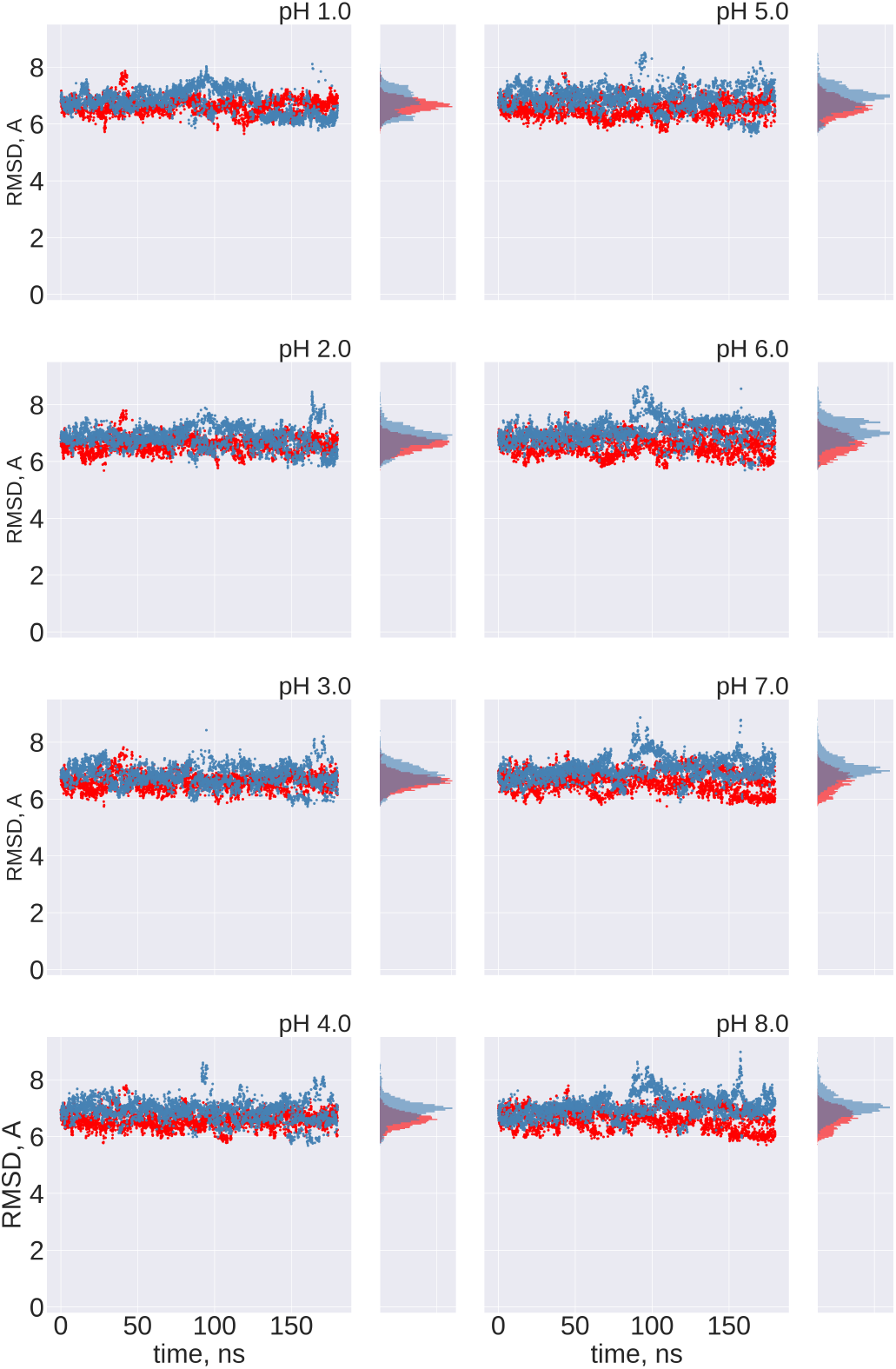
Cross-comparison of FPPR RMSD values as a metric for conformational transitions. FPPR RMSD values vs time are shown during two pH-REMD simulations initiated with different FPPR conformations. In each, the RMSD is calculated using the *other* FPPR conformation as the reference structure. Red symbols - initially extended, with backbone RMSD calculated relative to the experimental compact conformation; blue symbols - initially compact, with backbone RMSD calculated relative to the experimental extended conformation. Large values at all pH ranges indicate that no simulation samples a transition to the alternate FPPR conformation.

To confirm the results of the RMSD analysis, trajectory frames from both simulations across all pH ranges were combined and clustered with hierarchical agglomerative (bottom-up) approach using all non-hydrogen FPPR atoms (see Methods). The simulations starting from the two different FPPR conformations share no clusters, regardless of the pH (Figure S8). Frames from simulations starting in the extended conformation fall into a single cluster, while frames from the initially-compact conformation form two clusters, with a dominant cluster that is similar to the compact conformation. These observations confirm that no transition between conformations happens within 180 ns at any pH value.

Since both FPPR conformations appear to be kinetically trapped during the 180 ns pH-REMD simulations, we extended one of the three ph-REMD runs to 900 ns for both FPPR conformations. Even with the longer timescale, no transitions between conformations were sampled. This precludes the direct calculation of a pH-dependent equilibrium constant for the FPPR conformational change.

However, the lack of transitions between different FPPR conformations allows us to assign the pH-REMD titration profiles uniquely to a single FPPR conformation type. We therefore exploited the lack of conformational switching to compare the protonation behavior of the alternate FPPR conformations at each simulated pH, focusing on endosomal pH values of 4.0 to 6.5.

### Calculating pKa values for each FPPR conformation via pH-REMD

To test the hypothesis that the equilibrium distribution is pH-dependent, titration curves of titratable residues in the two FPPR conformations were compared. If the equilibrium distribution is indeed pH-dependent, the set of titration curves should differ.^38^ Here, the most important residues are those exhibiting *increased* pKa values compared to the reference pKa, since this would lead to protonation changes when moving from extracellular to endosomal pH. Downshifted pKa values would be less likely to affect the FPPR over the biologically-relevant pH range.

The titration curves of the majority of the 13 acidic side chains are similar between FPPR conformations, and the pKa values deviate by no more than 1 unit from the reference pKa values, as illustrated in Table 3 and Table S2, as well as Figure S9.

**Table 3:** pKa values of the 13 acidic side chains in the FPPR region, calculated from pH-REMD simulations. Only two have significantly different pKas in the different FPPR conformations (highlighted in red). Four residues have pKas shifted significantly away from the reference values, but the shifts are similar in the different conformations (indicated in italics). Error bars were obtained from independent simulations of each FPPR conformation type.

| Residue | Chain | Extended<br>pKa | Compact<br>pKa |
| --- | --- | --- | --- |
| Glu 281 | B | $3.8 \pm 0.0$ | $3.7 \pm 0.3$ |
| *Glu 554* | A | $3.0 \pm 0.1$ | $2.8 \pm 0.1$ |
| *Asp 568* | A | $1.4 \pm 0.1$ | $1.1 \pm 0.3$ |
| Asp 571 | A | $3.2 \pm 0.0$ | $3.8 \pm 0.2$ |
| *Asp 574* | A | *negative* | *negative* |
| Glu 583 | A | $3.6 \pm 0.0$ | $3.2 \pm 0.1$ |
| *Asp 586* | A | *negative* | *negative* |
| Asp 614 | A | $3.8 \pm 0.2$ | $3.7 \pm 0.4$ |
| Glu 619 | A | $4.7 \pm 0.0$ | $4.5 \pm 0.1$ |
| Asp 830 | B | $4.9 \pm 0.1$ | $5.4 \pm 0.5$ |
| **Asp 839** | B | **$1.6 \pm 0.1$** | **$2.9 \pm 0.2$** |
| Asp 843 | B | $4.4 \pm 0.3$ | $4.3 \pm 0.5$ |
| **Asp 848** | B | **$4.5 \pm 0.2$** | **$2.7 \pm 0.1$** |

The only amino acid whose pKa value exhibits an upward shift is Asp830. However, this shift occurs in both conformations, suggesting minimal contribution to a pH-dependent conformational switch. Other pKa values also are shifted downward from the reference values, but the shifts again are similar for compact and extended FPPR. For example, Asp574 and Asp586 demonstrate significant shifts in pKa values towards the negative range in both conformations. The pKa values of Asp568 and Glu554 show a downward shift as well. All four are located above the FPPR in the CTD1 domain, as shown in Figure 4.

Only Asp848 and Asp839 exhibit distinct titration curve patterns between compact and extended FPPR, as shown in Figure 8. In the compact conformation, Asp848 displays a downshifted pKa of 2.7 ± 0.1, while in the extended conformation, the pKa of 4.5 ± 0.2. Asp839 is downshifted in both FPPR conformations (2.9 ± 0.2 in the compact conformation, and even lower 1.6 ± 0.1 in the extended conformation) Figure 8.

**Figure 8:**
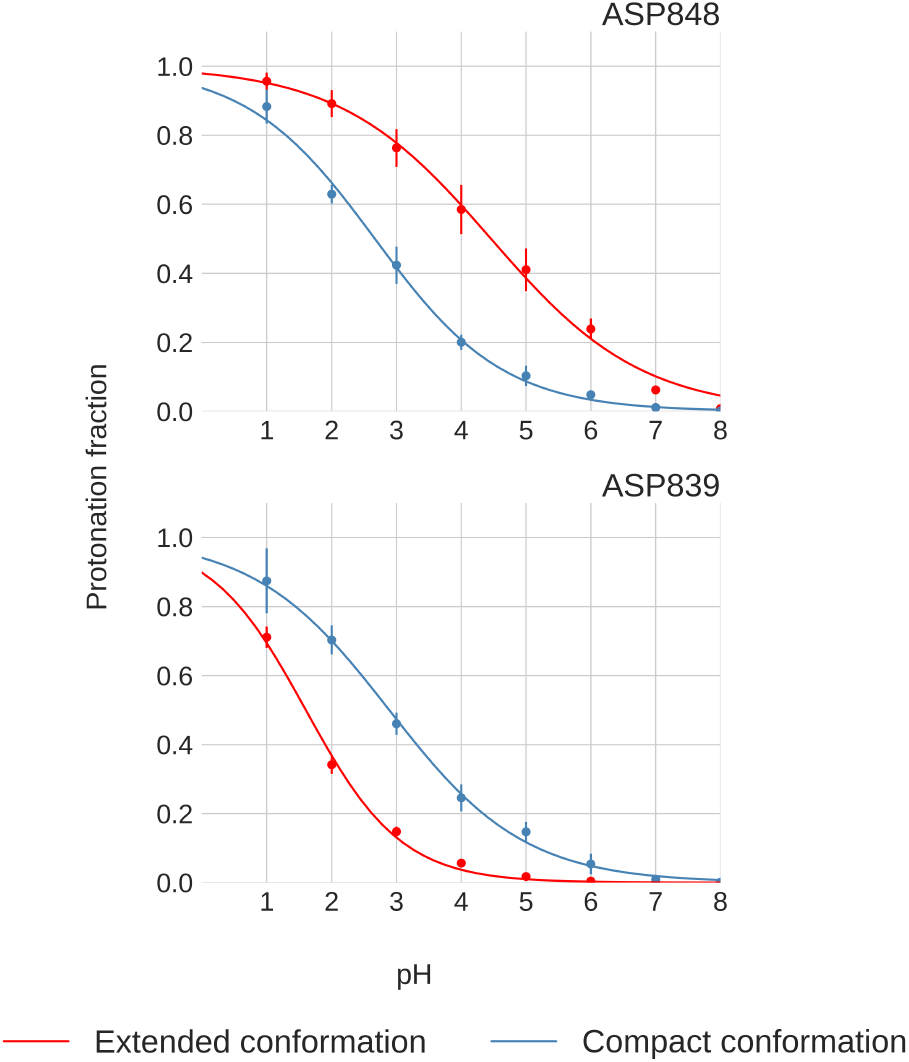
Titration curves of Asp848 and Asp839 from pH-REMD simulations. Error bars are calculated from independent simulatinos of that FPPR conformation

We next explored a structural basis for these calculated pKa shifts. In the compact conformation, the positioning of Asp848 at the N-terminal end of the helix (Ncap), as shown in Figure 4, may contribute to the observed lower pKa value due to interaction with the helix macrodipole.^53^ In contrast, in the extended conformation, Asp848 is not situated at the N-terminal of the helix, resulting in a pKa relatively unchanged from the reference value.

For Asp839, a downward shift in pKa occurs in both conformations, with a more pronounced shift in the extended conformation. The presence of nearby Lys835 as shown in Figure 4 may be the source of this shift, indicating a potential salt bridge formation. The reason for the magnified pKa shift in the extended conformation is less clear; however, since the pKa value is well below 4 and thus protonation is unlikely to play a biological role, we did not investigate further.

The presence of differences between individual pKa values across conformations suggest that the equilibrium distribution between compact and extended conformations might depend on pH. Despite these pH-dependent variations in specific residues,however, the total protonation of the 13 acidic side chains remains consistent across the two different conformations over the pH range of 4.0 to 6.5, as shown in Figure 9 and Table S3. This finding suggests that the total protonation for both FPPR conformations varies with pH, but is not influenced by conformation. This observation opposes the initial hypothesis proposing a pH-dependent equilibrium between FPPR conformations. MD simulations in the absence of experimental data should be treated with caution; however, these conclusions from our pH-REMD simulations are consistent with our PDB-based analysis of experimental FPPR conformations at different pH values.

**Figure 9:**
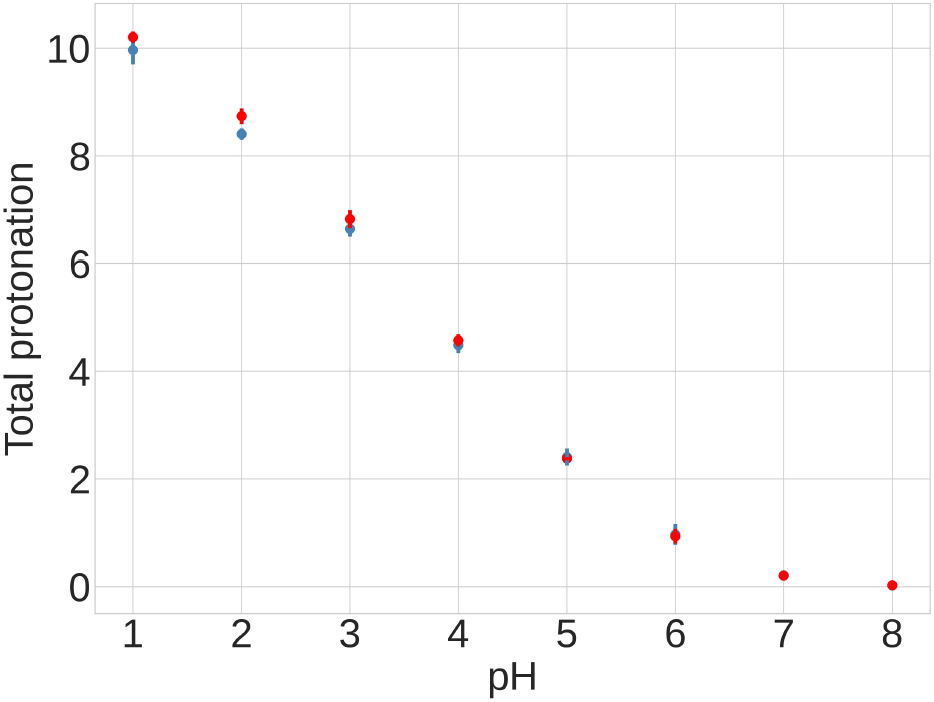
Results from pH-REMD simulations, showing the relationship between the total protonation of titratable groups and pH. The Y axis shows the average number of protonated acidic groups, with a maximum of 13 (total number of Asp and Glu titrated). Red - extended FPPR conformation, blue - compact FPPR conformation. Error bars are from independent simulations of the same FPPR conformation type.

### Forced competition between FPPR conformations at different pH values

The pH-REMD simulations described above involved initiation of all replicas in the same FPPR conformation (in the independent runs, either all extended, or all compact). No transitions to the other conformation were sampled, which permitted calculation of the pKa curves for the initial conformation as shown in Figure 8. However, inclusion of a single FPPR conformation type precluded observation of a preference for one conformation over another at a given pH. To overcome this obstacle, we initiated pH-REMD simulations in which both FPPR conformations were represented in the initial coordinates for a single pH-REMD run. After evolving the simulation for a period of time, the ratio of the two conformations is calculated at each pH. If one of the FPPR conformations is preferred at higher (or lower) pH than the other, we expected the REMD exchange process to lead to uneven sorting of conformations across the pH ladder, and a pH-dependent conformational preference to be apparent.

We initiated a 180 ns pH-REMD simulation with 8 replicas as before, but in this case the replicas initially at pH 1 to 4 started in the compact FPPR, while replicas at pH 5 to 8 started in the extended FPPR conformation. We analyzed the pH preference for each conformation by clustering the trajectory at each pH into two clusters using the k-means algorithm^12^ (see Table 4, Figure S10). The two conformations are similarly populated, within the uncertainty range, except at the extreme of pH 1. No trend is seen to favor either conformation as a function of pH. These data are consistent with the similar titration curves obtained from the separate pH-REMD calculations for each FPPR conformation. Overall, the simulations suggest a very similar free energy for each FPPR conformation at the biologically relevant pH range.

**Table 4:**
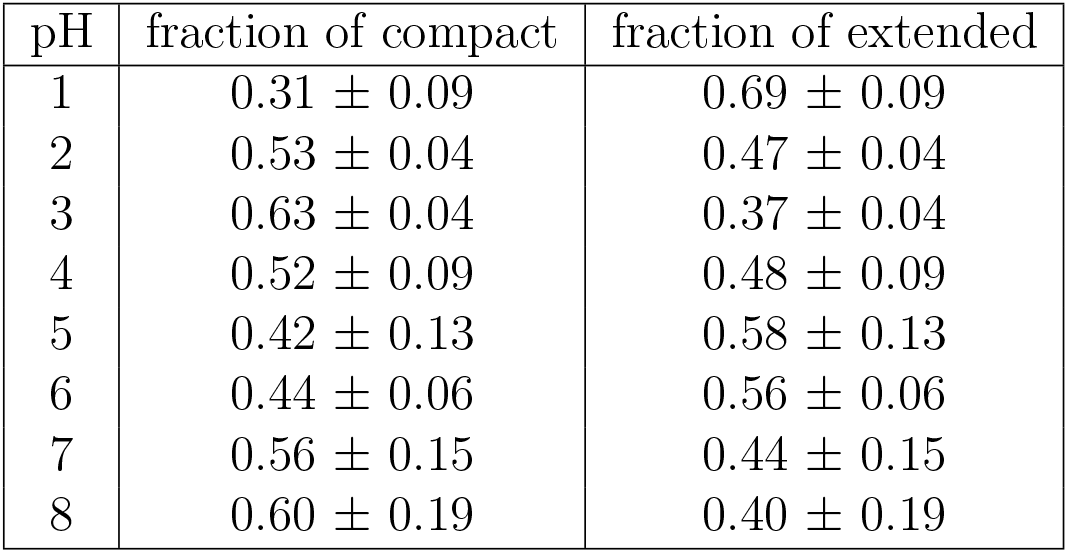
Data from pH-REMD with forced competition between different FPPR conformations. The table provides the fraction of simulation time spent in the compact and extended conformations at different pH values. Error bars were obtained from independent simulations.

| pH | fraction of compact | fraction of extended |
| --- | --- | --- |
| 1 | $0.31 \pm 0.09$ | $0.69 \pm 0.09$ |
| 2 | $0.53 \pm 0.04$ | $0.47 \pm 0.04$ |
| 3 | $0.63 \pm 0.04$ | $0.37 \pm 0.04$ |
| 4 | $0.52 \pm 0.09$ | $0.48 \pm 0.09$ |
| 5 | $0.42 \pm 0.13$ | $0.58 \pm 0.13$ |
| 6 | $0.44 \pm 0.06$ | $0.56 \pm 0.06$ |
| 7 | $0.56 \pm 0.15$ | $0.44 \pm 0.15$ |
| 8 | $0.60 \pm 0.19$ | $0.40 \pm 0.19$ |

## Conclusions

To test the hypothesis that the Fusion Peptide Proximal Region in SARS-CoV-2 spike glycoprotein exhibits pH dependence and controls pH-dependent RBD positioning, a comprehensive analysis of existing experimental structures was conducted, supplemented by new simulation data. Initial efforts focused on experimental structural analysis; the results did not support a difference in FPPR conformation as a function either of experimental pH or RBD position, although the scarcity of spike structures determined at low pH or with extended FPPR left remaining uncertainty. The extensive use of artificially stabilized spike constructs for structure determination may also be a complicating factor in analyzing spike PDB structures; very few structures correspond to a biologically-relevant spike sequence.

Calculating the pKa values of individual residues in selected experimental spike structures, with the aim of revealing distinct pKa values between conformations, yielded results that were poorly reproducible across different spike PDB structures with the same FPPR conformation type. pKa differences between different examples of the same FPPR conformation were comparable to the pKa differences between different FPPR conformations. The poor reproducibility of pKa values from static spike structures is likely attributable to the relatively low resolution of the cryo-EM structures, coupled with the sensitivity of pKa to local structure details and dynamics.

In order to estimate pKa values in the context of a dynamic ensemble, molecular dynamics simulations were conducted across a range of pH levels utilizing pH-REMD. Even after extending the pH-REMD to 1 microsecond, transitions between FPPR conformations were not observed at any pH due to the slow timescale of FPPR reorganization. This prevented the direct estimation of the free energy difference between conformations. However, it enabled the calculation of distinct pKa values for titratable side chains in the alternate FPPR conformations.

The pH-REMD simulations revealed divergent pKa values for two residues, Asp 848 and Asp 839, in different FPPR conformations. However, the total protonation showed no significant dependence on conformation, opposing the hypothesis of a pH-dependent equilibrium between conformations. Similarly, when pH-REMD simulations were initiated using both conformations in a single simulation, no preference was observed for either conformation as a function of pH.

In conclusion, the comprehensive analysis presented here, including meta-analysis of experimental conditions and structure correlations, as well as all-atom constant pH simulations, suggests that the FPPR is unlikely to be a pH-sensitive switch controlling the RBD position in the SARS-CoV-2 spike.

## Supporting information

6xlu.ff14SB_TIP3P.cpin

6xm0.ff14SB_TIP3P.cpin

AS4.ff14SB.off

GL4.ff14SB.off

frcmod.constph.ff14SB

## Acknowledgments

This work was supported by the Research Corporation for Science Advancement (COVID Initiative grant #27350). We gratefully acknowledge support from Henry and Marsha Laufer.

