## Supplemental Information for "Assessing pH-dependent Conformational Changes in the Fusion Peptide Proximal Region of the SARS-CoV-2 spike glycoprotein"

List S1: List of 700 SARS-CoV-2 full spike protein (Uniprot ID P0DTC2) cryo-EM structures with resolution 4 Å or better retrieved from the PDB on 08/3/2023.

6vsb, 6vxx, 6vyb, 6wps, 6wpt, 6x29, 6x2a, 6x2b, 6x2c, 6x6p, 6x79, 6xcm, 6xcn, 6xey, 6xf5, 6xf6, 6xkl, 6xlu, 6xm0, 6xm3, 6xm4, 6xm5, 6xr8, 6xra, 6xs6, 6z43, 6z97, 6zb4, 6zb5, 6zdh, 6zge, 6zgg, 6zgi, 6zhd, 6zow, 6zox, 6zoy, 6zoz, 6zp0, 6zp1, 6zp5, 6zp7, 6zwv, 6zxn, 7a25, 7a29, 7a4n, 7a94, 7ad1, 7akd, 7b18, 7bnm, 7bnn, 7byr, 7c2l, 7cab, 7cac, 7cai, 7cak, 7chh, 7cn4, 7ct5, 7cwl, 7cwm, 7cwn, 7cws, 7cwt, 7cwu, 7cyp, 7czp, 7czq, 7czr, 7czs, 7czt, 7czu, 7czv, 7czw, 7czx, 7czy, 7czz, 7d00, 7d03, 7d0b, 7d0c, 7d0d, 7ddd, 7df3, 7df4, 7dk4, 7dwy, 7dwz, 7dx0, 7dx1, 7dx2, 7dx3, 7dx5, 7dx6, 7dx7, 7dx8, 7dx9, 7dzw, 7dzx, 7dzy, 7e3k, 7e3l, 7e5r, 7e5s, 7e7b, 7e7d, 7e8c, 7e9n, 7e9o, 7e9q, 7eaz, 7eb0, 7eb3, 7eb4, 7eb5, 7edf, 7edg, 7edh, 7edi, 7edj, 7eh5, 7ej4, 7ej5, 7enf, 7epx, 7fae, 7faf, 7fb0, 7fb1, 7fb3, 7fb4, 7fcd, 7fce, 7fet, 7fjn, 7fjo, 7jji, 7jv4, 7jv6, 7jvc, 7jwb, 7jwy, 7jzl, 7jzn, 7k43, 7k4n, 7k8s, 7k8t, 7k8u, 7k8v, 7k8w, 7k8x, 7k8z, 7k90, 7k9h, 7k9j, 7kdg, 7kdh, 7kdi, 7kdj, 7kdk, 7kdl, 7ke4, 7ke6, 7ke7, 7ke8, 7ke9, 7kea, 7keb, 7kec, 7kj2, 7kj3, 7kj4, 7kj5, 7kkk, 7kkl, 7kml, 7kms, 7kmz, 7knb, 7kne, 7knh, 7kni, 7kqb, 7kqe, 7krq, 7krr, 7krs, 7ksg, 7l02, 7l06, 7l09, 7l2d, 7l2e, 7l2f, 7l3n, 7l56, 7l7e, 7l7k, 7laa, 7lab, 7lcn, 7ld1, 7ljr, 7lqv, 7lrt, 7ls9, 7lss, 7lwi, 7lwj, 7lwk, 7lwl, 7lwm, 7lwn, 7lwo, 7lwp, 7lwq, 7lws, 7lwt, 7lwu, 7lwv, 7lww, 7lxy, 7lxz, 7ly2, 7lyk, 7lyl, 7lym, 7lyn, 7lyo, 7lyq, 7m0j, 7m6e, 7m6f, 7m6g, 7m6h, 7m6i, 7mjg, 7mjh, 7mjj, 7mjk, 7mjm, 7mkl, 7mm0, 7mtc, 7mtd, 7mte, 7mw2, 7mw3, 7mw4, 7mw5, 7mw6, 7my2, 7my3, 7n0g, 7n0h, 7n1q, 7n1t, 7n1u, 7n1v, 7n1w, 7n1x, 7n5h, 7n8h, 7n9b, 7n9c, 7n9e, 7n9t, 7nd3, 7nd4, 7nd5, 7nd7, 7nd8, 7nd9, 7nda, 7ns6,

7nt9, 7nta, 7ntc, 7ny5, 7oan, 7od3, 7odl, 7p40, 7p77, 7p78, 7p79, 7p7b, 7q1z, 7q6e, 7q9f, 7q9g, 7q9j, 7q9m, 7qdg, 7qo7, 7qti, 7qur, 7qus, 7r13, 7r14, 7r16, 7r17, 7r18, 7r19, 7r1a, 7r1b, 7r40, 7r4i, 7r4q, 7r4r, 7r8m, 7r8n, 7r8o, 7ra8, 7rbv, 7rkv, 7ru1, 7ru2, 7ru3, 7ru5, 7rw2, 7s0c, 7s0d, 7s6i, 7s6j, 7s6k, 7s6l, 7sbk, 7sbl, 7sbp, 7sbq, 7sbs, 7sc1, 7sn3, 7so9, 7sob, 7soe, 7swx, 7sxr, 7sxs, 7sxt, 7sxu, 7sxv, 7sxw, 7sxx, 7sxz, 7sy1, 7sy3, 7sy5, 7sy7, 7t3m, 7t9j, 7t9k, 7tat, 7tb4, 7tb8, 7tca, 7tcc, 7tei, 7tex, 7tey, 7tf0, 7tf1, 7tf2, 7tf3, 7tf4, 7tf5, 7tgw, 7tgx, 7tgy, 7thk, 7tht, 7tla, 7tlb, 7tlc, 7tld, 7tm0, 7tnw, 7to4, 7tou, 7tov, 7tox, 7toy, 7tp0, 7tp1, 7tp2, 7tp7, 7tp8, 7tp9, 7tpa, 7tpc, 7tpe, 7tpf, 7tph, 7tpl, 7tpr, 7tyz, 7u0p, 7u0q, 7u0x, 7uap, 7uar, 7ub0, 7ub5, 7ub6, 7uhc, 7upw, 7upy, 7uz4, 7uz5, 7uz6, 7uz7, 7uz8, 7uz9, 7uza, 7v20, 7v23, 7v26, 7v2a, 7v76, 7v77, 7v78, 7v79, 7v7a, 7v7d, 7v7e, 7v7f, 7v7g, 7v7h, 7v7i, 7v7j, 7v7n, 7v7o, 7v7p, 7v7q, 7v7r, 7v7s, 7v7t, 7v7u, 7v7v, 7v7z, 7v81, 7v82, 7v83, 7v85, 7v86, 7v88, 7v89, 7v8a, 7v8c, 7vnc, 7vnd, 7vne, 7vq0, 7vrw, 7vx1, 7vx9, 7vxa, 7vxb, 7vxc, 7vxd, 7vxe, 7vxf, 7vxi, 7vxk, 7vxm, 7w92, 7w94, 7w98, 7w99, 7w9b, 7w9c, 7w9e, 7wbh, 7wcd, 7wcz, 7wd0, 7wd7, 7wd9, 7wdf, 7we7, 7we8, 7we9, 7wea, 7web, 7wec, 7wev, 7wg7, 7wg9, 7wgb, 7wgv, 7wgx, 7wgy, 7whb, 7whd, 7whi, 7whj, 7whk, 7wjy, 7wjz, 7wk2, 7wk3, 7wk4, 7wk5, 7wk9, 7wka, 7wly, 7wlz, 7wo5, 7woa, 7wob, 7woq, 7wor, 7wos, 7wou, 7wov, 7wp9, 7wpa, 7wpd, 7wpe, 7wpf, 7wqv, 7wrh, 7ws0, 7ws1, 7ws3, 7ws4, 7ws5, 7ws8, 7ws9, 7wtf, 7wti, 7wtk, 7wvn, 7wvo, 7wvp, 7wwi, 7wwj, 7wwl, 7wwm, 7wz1, 7wz2, 7x08, 7x6a, 7xch, 7xco, 7xdb, 7xdk, 7xdl, 7xic, 7xid, 7xiw, 7xix, 7xiy, 7xmx, 7xmz, 7xnq, 7xnr, 7xns, 7xo4, 7xo5, 7xo7, 7xo8, 7xoa, 7xob, 7xod, 7xst, 7xu1, 7y7j, 7y9s, 7y9z, 7ya0, 7yeg, 7yqt, 7yqu, 7yqv, 7yqw, 7yqx, 7yqy, 7yqz, 7yr1, 7yr2, 7yr3, 7z3z, 7z6v, 7z7x, 7z85, 7z86, 7z9q, 7zce, 7zr7, 7zr9, 7zrc, 7zrv, 7zss, 8csa, 8cxn, 8cxq, 8cy6, 8cy7, 8cy9, 8cya, 8cyb, 8cyc, 8cyd, 8df5, 8dli, 8dlj, 8dll, 8dlm, 8dlo, 8dlp, 8dlt, 8dlu, 8dlw, 8dlx, 8dlz, 8dm1, 8dm3, 8dm5, 8dm7, 8dm9, 8dxs, 8dzh, 8dzi, 8err, 8gs6, 8hc4, 8hhx, 8hhy, 7x93, 7xj6, 7xj8, 7zjl, 8a94, 8a99, 8c1v, 8d0z, 8f0g, 8fez, 8fu7, 8fu8, 8fu9, 8gjm, 8h00, 8h01, 8h3d, 8h3e, 8h3m, 8h3n, 8heb, 8hec, 8itu, 7wch, 7wcp, 7wt7, 7wt8, 7xd2, 7y1y, 7y1z, 7y20, 7y21, 7y71, 7ybh, 7ybj, 7ybl, 7ybm, 7yc5, 7yh7, 7yve, 7yvg, 7yvi, 7yvk, 7yvn, 7yvo, 7yvp, 8a95, 8aja, 8ajl, 8bon, 8cim, 8csj, 8d55, 8d56, 8d5a, 8dt8, 8elj, 8gou, 8gto, 8gtp, 8gtq, 8h07, 8h08, 8ios, 8iot, 8iou

Table S1: pKa values of 13 titratable residues of a Spike fragment calculated using PROPKA3 using PDB files in both FPPR conformations.

| | | Extended conformation | | Compact conformation | |
|---|---|---|---|---|---|
| Residue | Chain | 6XM0 | 7LQV | 6XLU | 7WGX |
| Glu 281 | B | 3.6 | 4.0 | 4.1 | 4.7 |
| Glu 554 | A | 4.2 | 3.9 | 4.8 | 3.9 |
| Asp 568 | A | 4.7 | 4.9 | 5.0 | 7.2 |
| Asp 571 | A | 4.3 | 4.3 | 4.4 | 4.4 |
| Asp 574 | A | 6.0 | 6.8 | 6.7 | 4.2 |
| Glu 583 | A | 4.8 | 5.2 | 4.6 | 3.2 |
| Asp 586 | A | 4.7 | 5.0 | 5.0 | 4.9 |
| Asp 614 | A | 5.4 | 5.7 | 7.0 | 4.8 |
| Glu 619 | A | 4.7 | 4.8 | 4.1 | 3.5 |
| Asp 830 | B | 4.0 | 4.5 | 4.5 | 4.0 |
| Asp 839 | B | 3.9 | 3.8 | 3.6 | 4.4 |
| Asp 843 | B | 4.2 | 4.3 | 6.0 | 3.8 |
| Asp 848 | B | 4.3 | 4.4 | 4.9 | 3.8 |

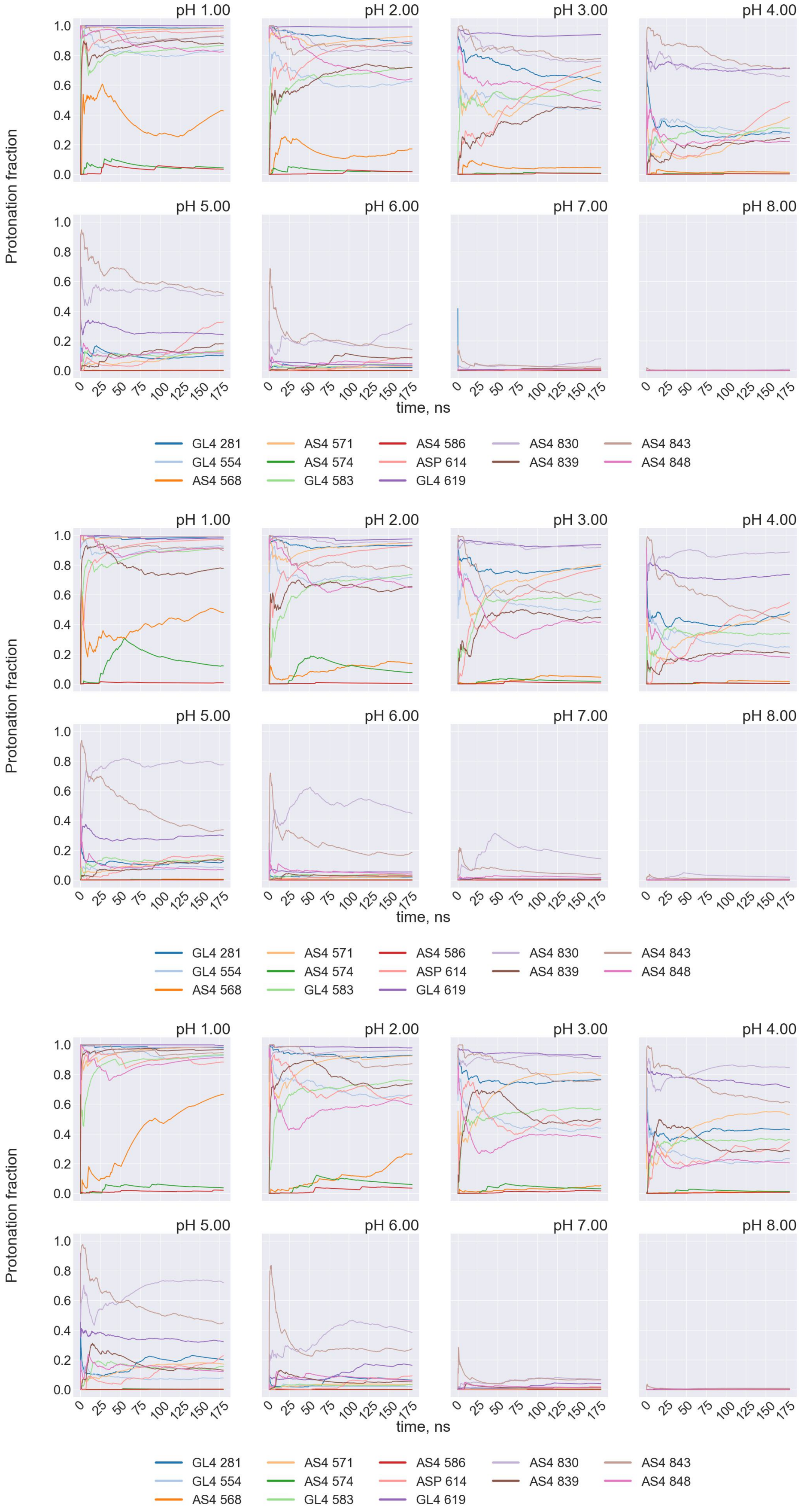


Figure S1: Time series of the cumulative protonated fraction of 13 titratable residues in initially compact conformation in pH-REMD. Simulations were repeated in triplicate.

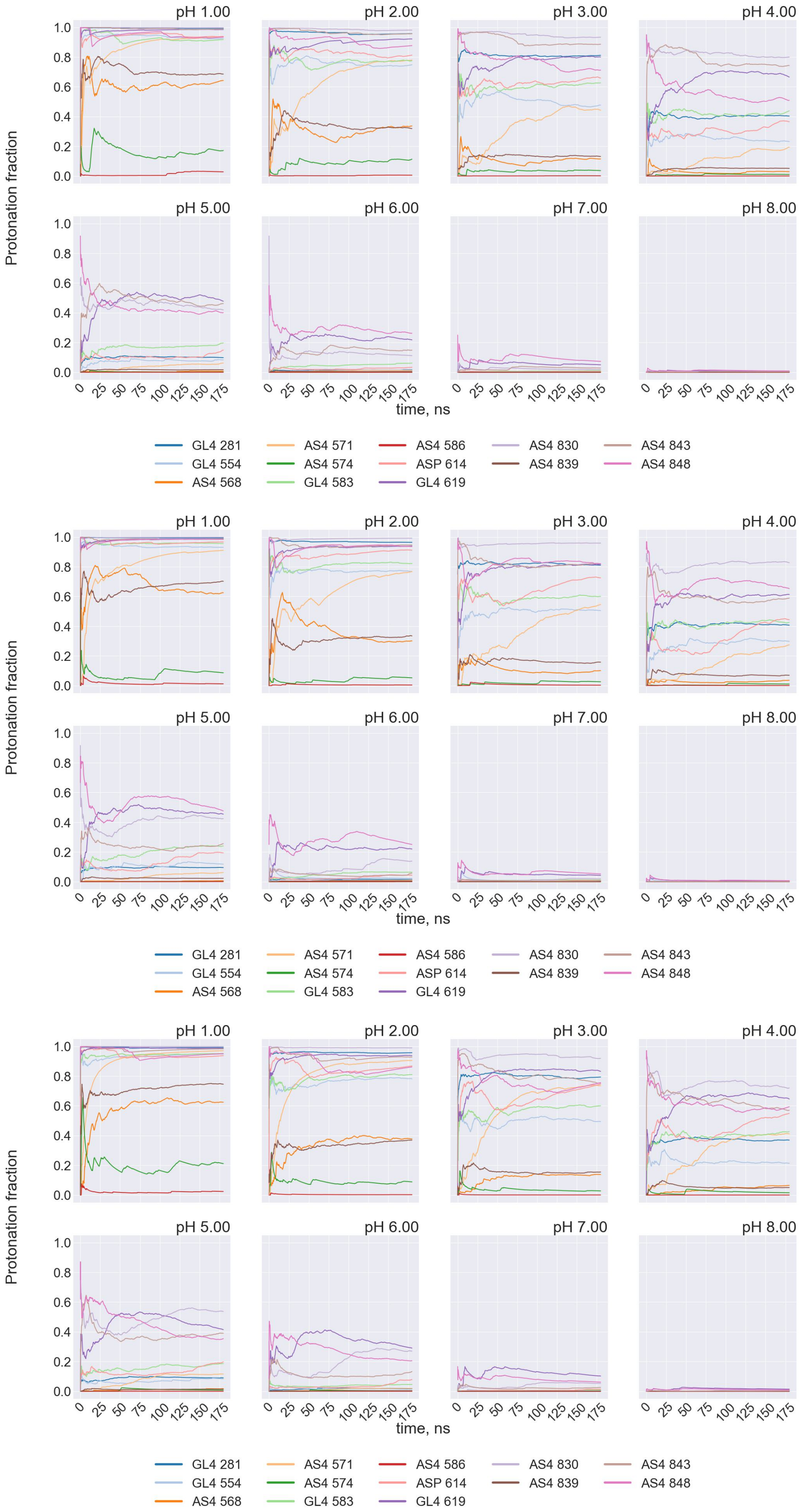


Figure S2: Time series of the cumulative protonated fraction of 13 titratable residues in initially extended conformation in pH-REMD. Simulations were repeated in triplicate..

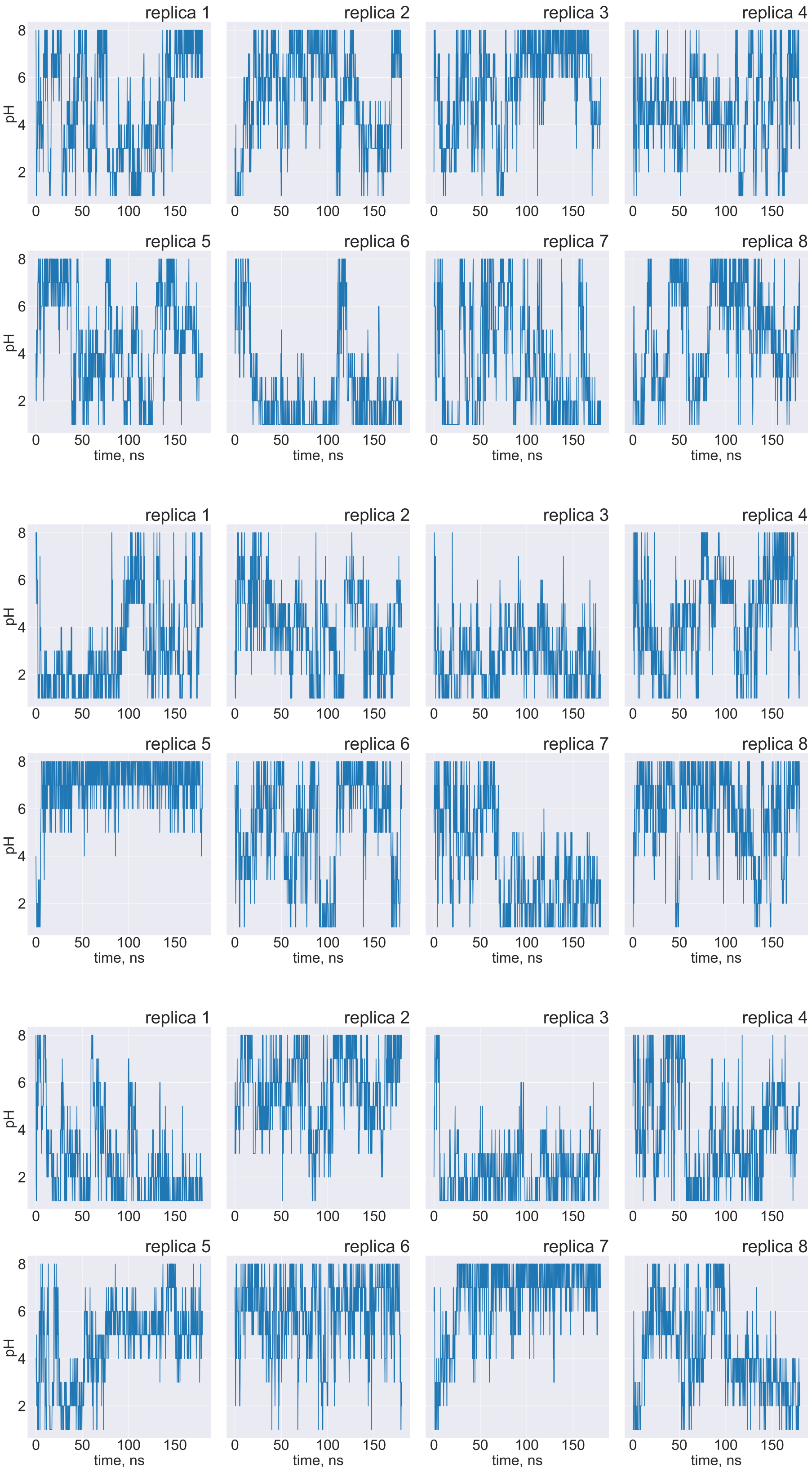


Figure S3: Time series of the pH for individual replica trajectories in pH-REMD in extended conformation. Simulations were repeated in triplicate.

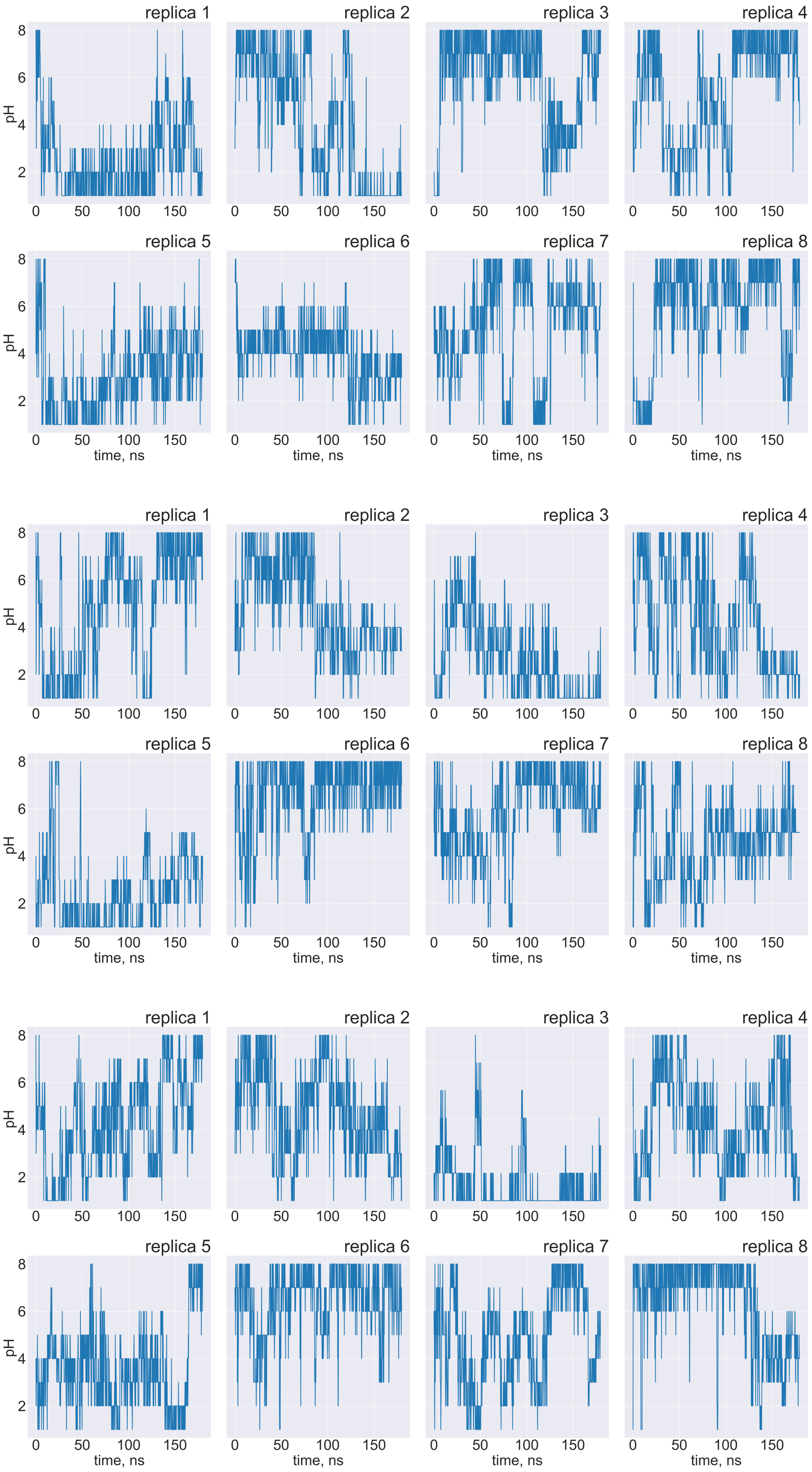


Figure S4: Time series of the pH for individual replica trajectories in pH-REMD in compact conformation. Simulations were repeated in triplicate.

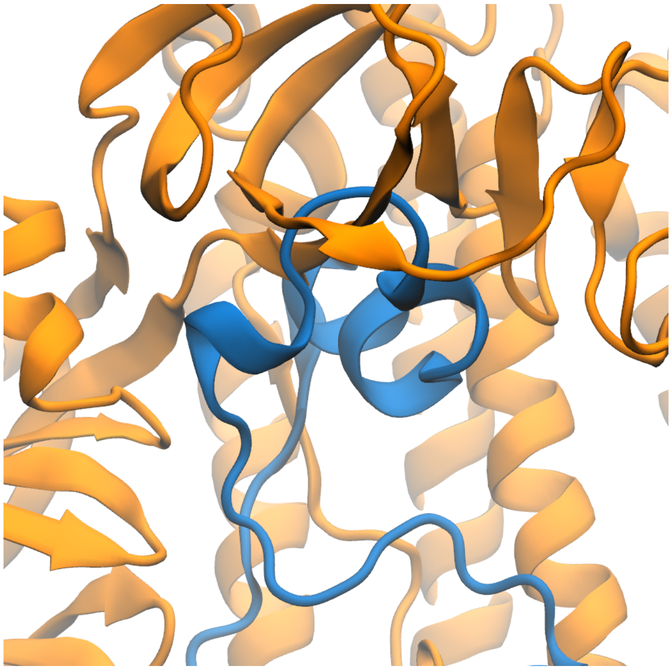

Figure S5: FPPR in compact conformation in PDB 7DZW. FPPR (chain C) is shown with blue ribbon and other regions use orange. The FPPR has significant clash with the CTD1 domain above it.

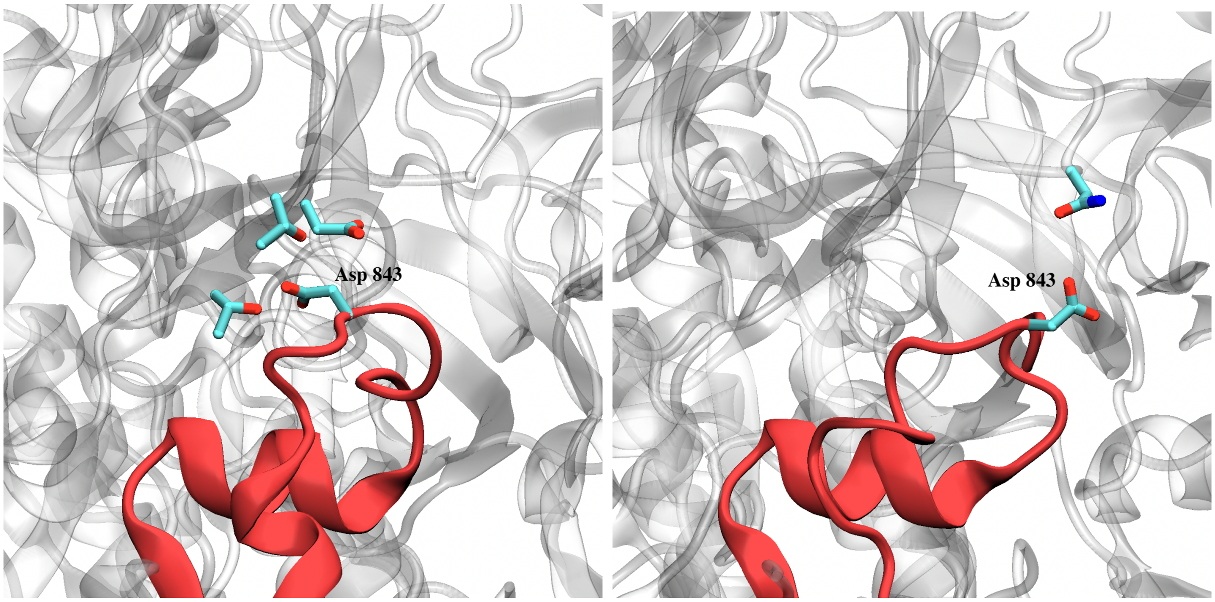


Figure S6: FPPR in compact conformation in PDB 6XLU (left) and 7WGX (right). FPPR (chain B) is shown with red ribbon and other regions use gray. In 6XLU Asp843 is surrounded by Asp586, Thr553 and Thr588 and has a pKa of 6.0. In 7WGX Asp843 is surrounded by Asn556 and has a pKa of 3.8.

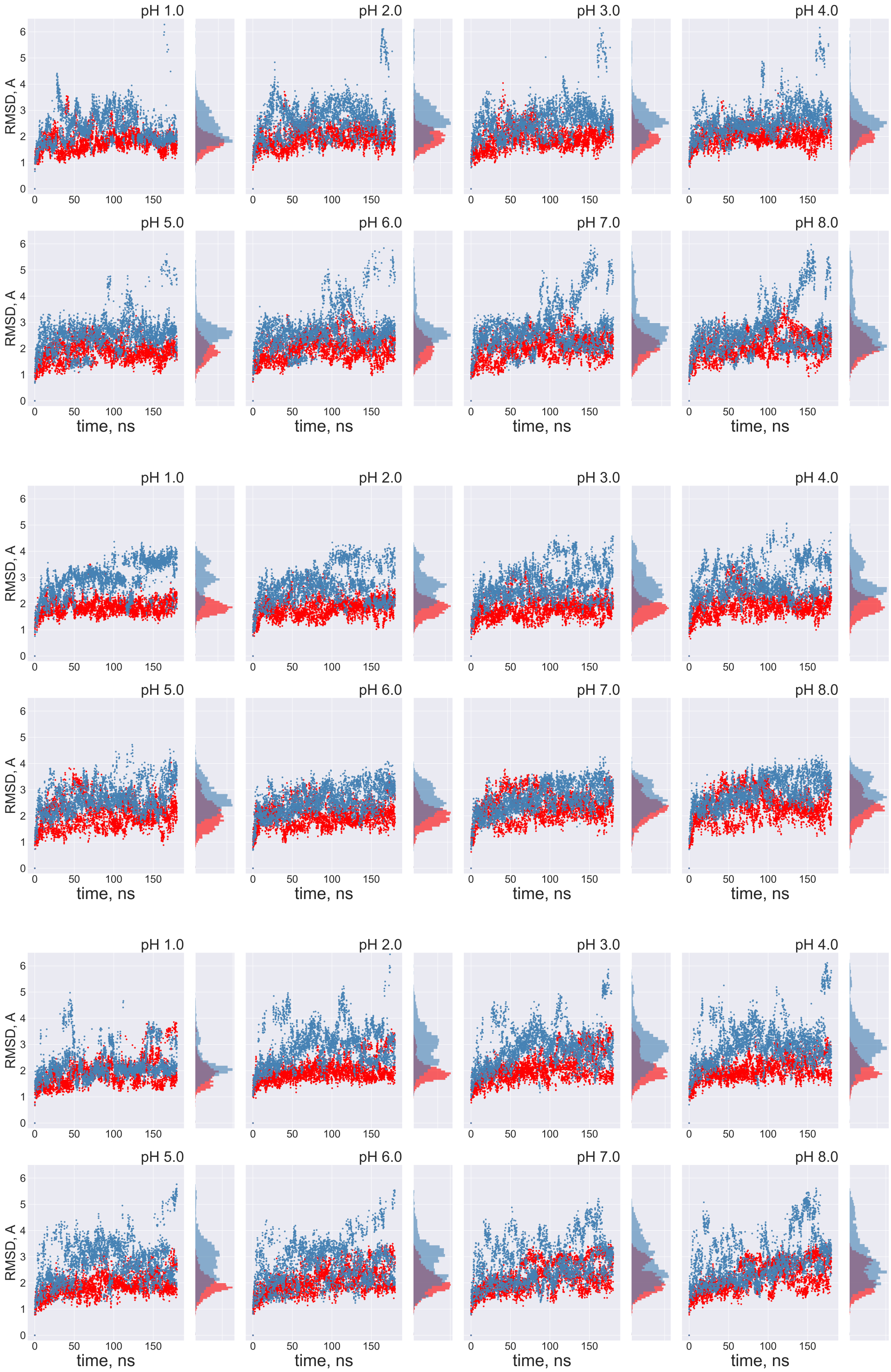


Figure S7: Time series of the RMSD for FPPR backbone atoms with a histogram on the right. Data are shown at different pH values during two pH-REMD simulations, each initiated with a different FPPR conformation. Red symbols - initially extended simulation, with backbone RMSD calculated using the experimental extended conformation; blue symbols - initially compact simulation, with backbone RMSD calculated using to the experimental compact conformation. Low values indicate that the initial FPPR conformation is stable at all pH values. 3 repeats

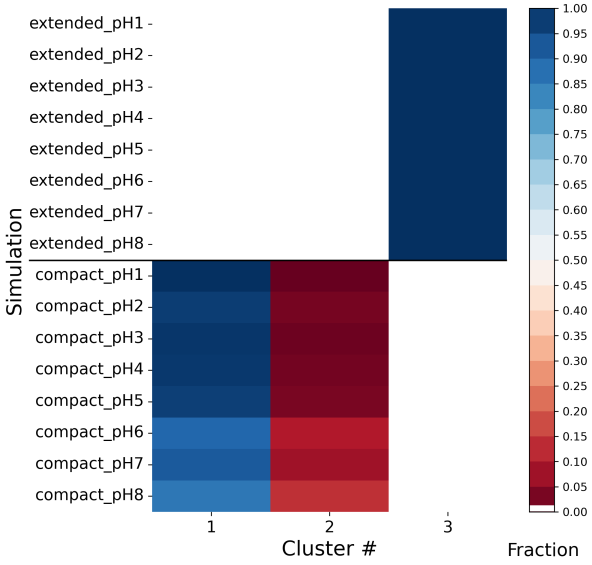


Figure S8: Trajectory frames from both simulations across all pH ranges were combined and clustered with hierarchical clustering using symmetry-corrected RMSD for all non-hydrogen FPPR atoms. The fractional population for each cluster is color-coded. Each row sums to 1.0. Frames from simulations starting in the extended conformation fall into a single cluster, while frames from the initially-compact conformation form two clusters.

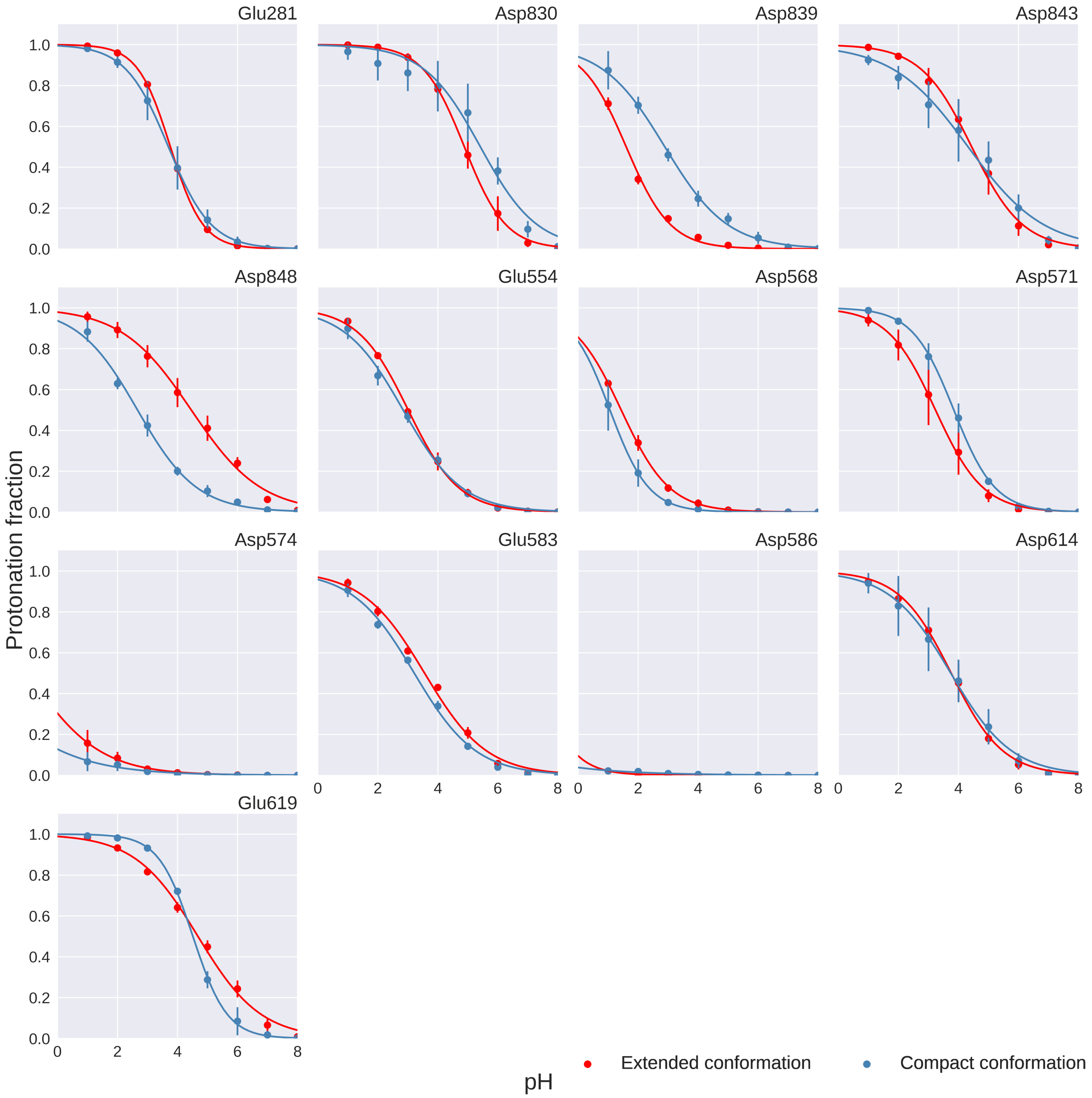


Figure S9: Titration curves of 13 titratable residues in two different FPPR conformations from pH-REMD simulations. Red - extended conformation, blue - compact conformation. Error bars are obtained from the independent simulations.

Table S2: Mean pKa values and Hill coefficients with standard deviation for 13 titratable residues in both conformations from 3 repeats of pH-REMD simulations. Red highlights 2 residues that have different pKas between the two conformations.

| | extended conformation | | compact conformation | |
|---|---|---|---|---|
| residue | pKa | Hill coefficient | pKa | Hill coefficient |
| Glu 281 | 3.8 ± 0.0 | 0.8 | 3.7 ± 0.3 | 0.6 |
| Glu 554 | 3.0 ± 0.1 | 0.5 | 2.8 ± 0.1 | 0.4 |
| Asp 568 | 1.4 ± 0.1 | 0.5 | 1.1 ± 0.3 | 0.7 |
| Asp 571 | 3.2 ± 0.0 | 0.5 | 3.8 ± 0.2 | 0.6 |
| Asp 574 | negative | - | negative | - |
| Glu 583 | 3.6 ± 0.0 | 0.4 | 3.2 ± 0.1 | 0.4 |
| Asp 586 | negative | - | negative | - |
| Asp 614 | 3.8 ± 0.2 | 0.5 | 3.7 ± 0.4 | 0.4 |
| Glu 619 | 4.7 ± 0.0 | 0.4 | 4.5 ± 0.1 | 0.8 |
| Asp 830 | 4.9 ± 0.1 | 0.6 | 5.4 ± 0.5 | 0.5 |
| Asp 839 | 1.6 ± 0.1 | 0.6 | 2.9 ± 0.2 | 0.4 |
| Asp 843 | 4.4 ± 0.3 | 0.5 | 4.3 ± 0.5 | 0.3 |
| Asp 848 | 4.5 ± 0.2 | 0.4 | 2.7 ± 0.1 | 0.4 |

Table S3: Average protonation of 13 titratable residues in 3 repeats of pH-REMD simulations of compact and extended conformations.

| | compact conformation | | | | extended conformation | | | |
|---|---|---|---|---|---|---|---|---|
| pH | 1 | 2 | 3 | mean ± std | 1 | 2 | 3 | mean ± std |
| 1 | 9.7 | 10.0 | 10.2 | 10.0 ± 0.3 | 10.1 | 10.1 | 10.3 | 10.2 ± 0.1 |
| 2 | 8.3 | 8.5 | 8.4 | 8.4 ± 0.1 | 8.6 | 8.7 | 8.9 | 8.7 ± 0.1 |
| 3 | 6.5 | 6.8 | 6.6 | 6.6 ± 0.1 | 6.6 | 6.9 | 6.9 | 6.8 ± 0.2 |
| 4 | 4.3 | 4.5 | 4.6 | 4.5 ± 0.1 | 4.4 | 4.6 | 4.6 | 4.6 ± 0.1 |
| 5 | 2.4 | 2.3 | 2.6 | 2.4 ± 0.2 | 2.4 | 2.3 | 2.4 | 2.4 ± 0.0 |
| 6 | 0.8 | 0.9 | 1.2 | 1.0 ± 0.2 | 0.9 | 0.8 | 1.1 | 0.9 ± 0.1 |
| 7 | 0.2 | 0.2 | 0.2 | 0.2 ± 0.0 | 0.2 | 0.1 | 0.3 | 0.2 ± 0.1 |
| 8 | 0.0 | 0.0 | 0.0 | 0.0 ± 0.0 | 0.0 | 0.0 | 0.0 | 0.0 ± 0.0 |

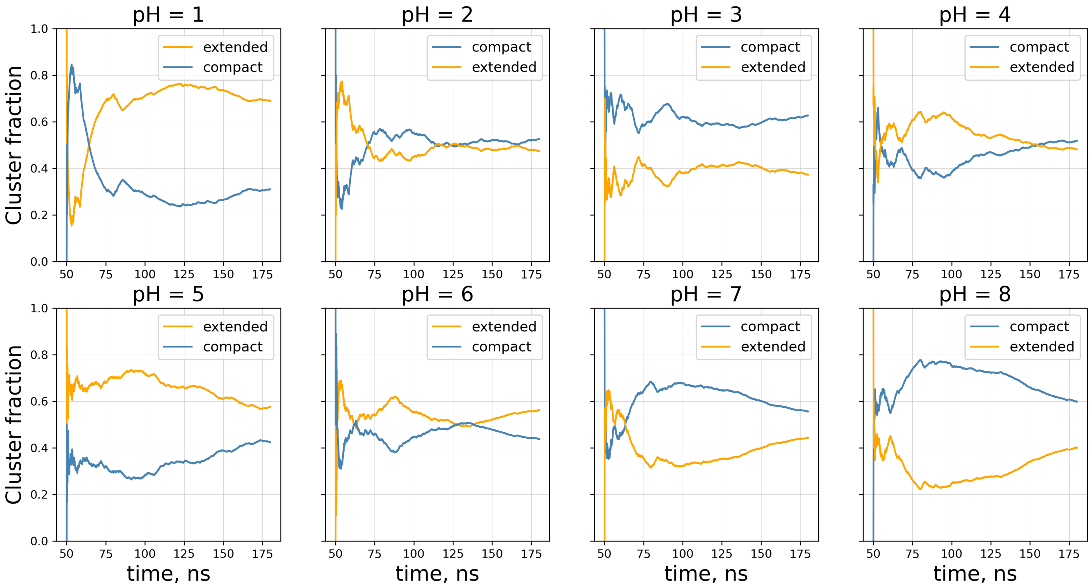


Figure S10: Time series of the fraction of each conformation at different pH values along the pH-REMD simulation started with replicas of both conformations.
